# Alternative splicing of a potato disease resistance gene maintains homeostasis between development and immunity

**DOI:** 10.1101/2023.06.09.544375

**Authors:** Biying Sun, Jie Huang, Liang Kong, Chuyun Gao, Fei Zhao, Jiayong Shen, Tian Wang, Kangping Li, Luyao Wang, Yuanchao Wang, Dennis A. Halterman, Suomeng Dong

## Abstract

Plants possess a robust and sophisticated innate immune system against pathogens. The intracellular receptors with nucleotide-binding, leucine-rich repeat (NLR) motifs recognize pathogen-derived effector proteins to trigger the immune response. To balance plant growth and rapid pathogen detection, NLR expression is precisely controlled in multifaceted ways. The alternative splicing (AS) of introns in response to infection is recurrently observed but poorly understood. Here we report that the potato NLR gene *RB* undergoes AS of its intron, resulting in two transcriptional isoforms, which coordinately regulate plant immunity and growth homeostasis. During normal growth, *RB* predominantly exists as intron-retained isoform *RB_IR*, encoding a truncated protein containing only the N-terminus of the NLR. Upon late blight infection, the pathogen induces intron splicing of *RB*, increasing the abundance of *RB_CDS*, which encodes a full-length and active R protein. By deploying the *RB* splicing isoforms fused with a *luciferase* reporter system, we identified IPI-O1 (also known as Avrblb1), the RB cognate effector, as a facilitator of *RB* AS. IPI-O1 directly interacts with potato splicing factor StCWC15, resulting in altered localization of StCWC15 from the nucleoplasm to the nucleolus and nuclear speckles. Mutations in IPI-O1 that eliminate StCWC15 binding also disrupt StCWC15 re-localization and *RB* intron splicing. Thus, our study reveals that StCWC15 serves as a surveillance facilitator sensing the pathogen-secreted effector, and regulates the trade-off between *RB*-mediated plant immunity and growth, expanding our understanding of molecular plant-microbe interactions.

**One-sentence summary:** Potato resistance gene *RB* balances plant growth and immunity through AS (alternative splicing), while pathogen-secreted effector IPI-O1 mediates AS of *RB* by targeting the conserved splicing factor StCWC15, further increasing the *RB_CDS* expression level to activate immunity.

## Introduction

Upon pathogen infection, plants have evolved two major mechanisms for recognition of pathogen effectors and elicitation of plant defense (Jones and Dangl, 2006). One utilizes plasma membrane-localized pattern-recognition receptors (PRRs) to perceive pathogen-associated molecular patterns (PAMPs), leading to PAMP-triggered immunity (PTI) (Couto and Zipfel, 2016; Yu et al., 2017; Zhou and Zhang, 2020). The other uses intracellular NLR immune receptors to detect cytoplasmic effectors introduced by the pathogen, thereby activating effector-triggered immunity (ETI; Jones and Dangl, 2006; Dodds and Rathjen, 2010; Cui et al., 2015). During ETI, NLRs oligomerize to form “resistosome” structures (Wang et al., 2019; Ofir et al., 2021) and the response culminates in programmed cell death, known as the hypersensitive response (HR), to prevent pathogen spread. The enormous diversity of *NLR* genes in plants demonstrates their important role in plant immunity (Li et al., 2015; Van de Weyer et al., 2019).

Late blight, caused by oomycete *Phytophthora infestans*, is the most devastating disease in potato production (Haverkort et al., 2009; Kamoun et al., 2015; Sun et al., 2022). Deployment of host resistance (*R*) genes is a crucial strategy for controlling the disease and many potato cultivars have been bred to introduce resistance genes against *P. infestans* (*Rpi* genes) that originate from potato wild relatives (Kamoun and Smart, 2005; Karki et al., 2021). More than 70 *Rpi* genes have been identified and mapped in 32 *Solanum* species (Paluchowska et al., 2022). Among these *R* genes, *RB* was cloned from the wild diploid potato species *Solanum bulbocastanum*, which is known for its durable late blight resistance (Song et al., 2003; van der Vossen et al., 2003). Like most plant *R* genes, *RB* encodes an NLR receptor which recognizes *P. infestans* effector IPI-O1 (also known as Avrblb1) to activate immune responses (Vleeshouwers et al., 2008; Champouret et al., 2009). Previous studies hypothesized that IPI-O1 might change the conformation of RB to facilitate the formation a functional multimeric complex, thereby activating *RB*-mediated disease resistance (Chen et al., 2012; Zhao and Song, 2021). Consequently, for *RB* with a durable-resistance properties, the functions of *RB*-mediated plant immunity need to be further determined. Despite their usefulness in activating resistance, constitutive expression of *NLR* genes causes auto-immunity, leading to a negative effect on plant growth and development (Cheng et al., 2011; Deng et al., 2019; Cui et al., 2020). Therefore, the expression of *NLR* genes needs to be precisely regulated to maintain homeostasis between growth and immunity. The regulatory mechanisms for the expression of NLRs are mainly via transcription regulation, RNA processing, post-translational modification and proteasome-mediated degradation (Li et al., 2015). Intron splicing is common in post-transcriptional RNA processing and is executed by a supramolecular ribonucleoprotein (RNP) complex known as the spliceosome (Reddy, 2007).

Intron retention during RNA processing is a form of alternative splicing (AS) and can lead to changes in the translated protein. AS contributes to transcriptome and proteome diversity in animals and plants and it has been shown to play an important role in regulating plant immunity (Reddy et al., 2013; Wu et al., 2016; Bazin et al., 2020; Zhang et al., 2022). Several *NLR* genes have been demonstrated to undergo AS, suggesting that this may be an important mechanism for regulation of this important class of immunity-related genes (Dinesh-Kumar and Baker, 2000; Halterman et al., 2003; Halterman and Wise, 2006; Xu et al., 2012; Sanchez-Martin et al., 2021). However, the molecular mechanisms that control *NLR* gene AS and the significance of AS in maintaining plant growth and immunity homeostasis is incompletely understood.

In this work, we report that the potato resistance gene *RB* undergoes alternative splicing, resulting in generation of two transcripts *RB_IR* and *RB_CDS.* Removal of the intron results in intact RB protein (RB_CDS) that confers resistance to *P. infestans*, while intron retention results in a truncated RB protein (RB_IR) that cannot protect against the pathogen. Upon late blight infection, *P. infestans* is able to modulate the AS of *RB*, increasing the abundance of *RB_CDS* to activate *RB*-mediated immunity. Interaction of *P. infestans* effector IPI-O1 with potato splicing factor CWC15 in the nucleus leads to an increase of *RB* intron splicing during infection. This process balances plant growth and defense and provides a unique molecular basis for NLR recognition of pathogen effectors.

## Results

### The AS of *RB* is specifically modulated by *P. infestans* infection

The *RB* gene encompasses a single 679-nucleotide (nt) intron initiating 428 nt downstream of the translation start codon and terminating at 1107 nt (Fig. 1A). We performed rapid amplification of cDNA ends (5’ RACE) using total RNA from non-inoculated *RB* transgenic potato cv. Desiree expressing a single genomic copy of *RB* with its native promoter (Zhu et al., 2015), and the transcriptional isoforms were identified that correspond to the transcripts with the intron spliced (*RB_CDS*) or retained (*RB-IR*) (Fig. 1A). The presence of these two isoforms was validated using RT-PCR of *RB* transgenic potato and *Nicotiana benthamiana*. Sequencing of the amplified products confirmed the presence of *RB_IR* and *RB_CDS* transcripts. (Fig. 1B and Supplemental Fig. S1A). Furthermore, the two transcripts caused by alternative intron splicing were also detected in RNA from the diploid potato species *Solanum bulbocastanum* clone PT29, where *RB* was identified (Fig. 1B) (Song et al., 2003). *Rpi-sto1*, an *RB*-homologous gene from *S. stoloniferum*, also showed the same two transcripts detected by RT-PCR in *Rpi-sto1* transgenic potato (Fig. 1B) (van der Vossen et al., 2003; Wang et al., 2008). Splicing of introns in pre-mRNA is directed by spliceosome targeting of particular sequences at the intron/exon junctions called splice sites (Wan et al., 2020), which are commonly found as a conserved GT dinucleotide at the 5’ splice site and a conserved AG dinucleotide at the 3’ splice site (Sheth et al., 2006). We generated *RB* splicing mutant *RB_IVS* (IVS=intervening sequence), with synonymously mutated splice sites in *RB* from GT to GA (5’) and from AG to CG (3’) (Fig. 1A), and transformed it into potato and *N. benthamiana*. We found that *RB_IVS* is unable to undergo intron splicing in both transgenic potato and *N. benthamiana* (Fig. 1B). Among the two transcripts of *RB*, *RB_CDS* is predicted to encode the entire NLR protein, while *RB_IR* encodes a truncated protein containing only the N-terminus of the NLR protein, due to a premature termination codon (PTC) in the intron (Fig. 1A). As expected, immunoblot analysis showed that RB fused with green florescent protein at the N-terminus (GFP-RB) produced two bands corresponding to RB_CDS and RB_IR, respectively (Fig. 1C). Together, the data support that *RB* undergoes AS to produce two different transcriptional isoforms that each encode different size proteins.

**Figure 1.**
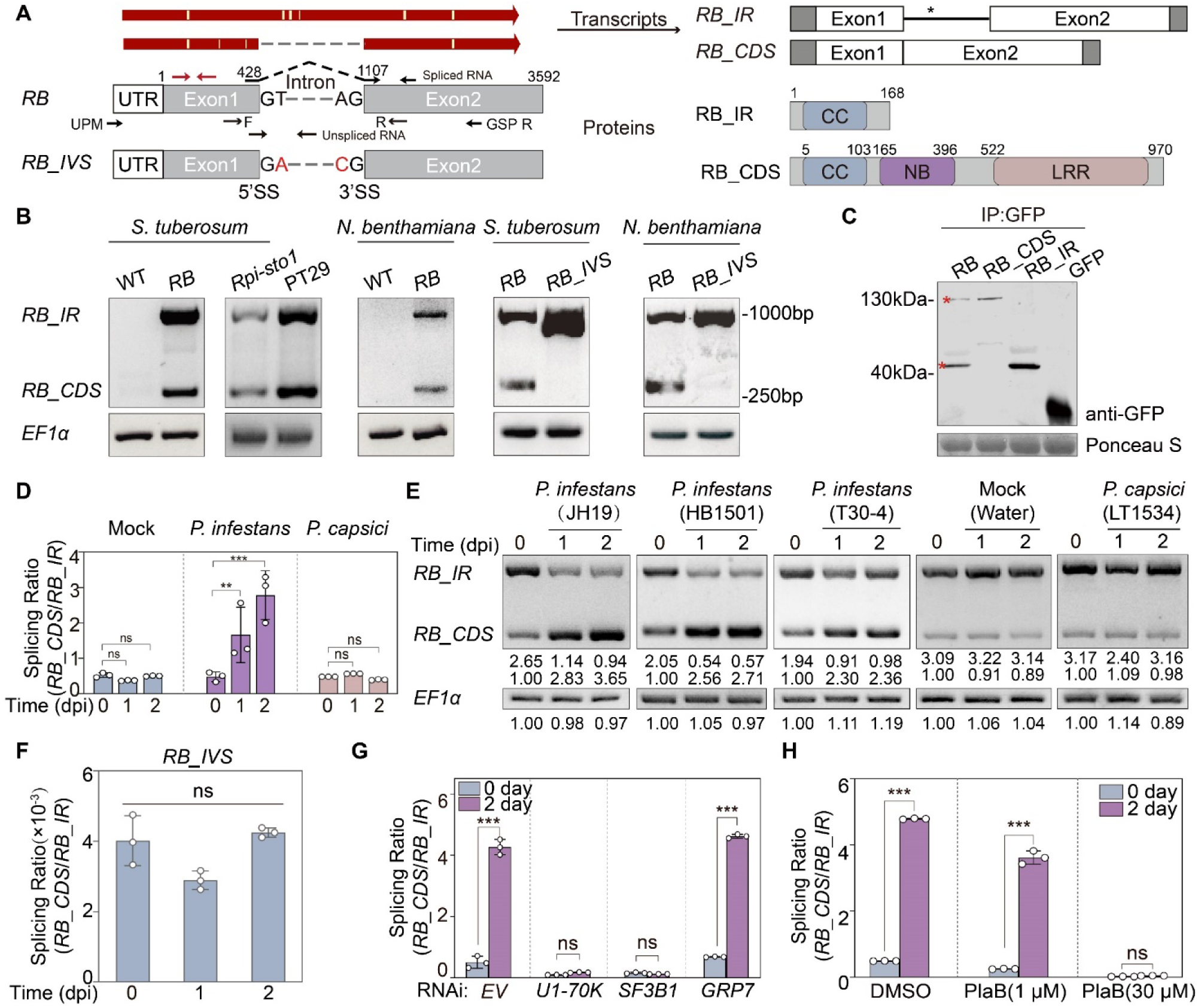
The AS of *RB* is specifically modulated by *P. infestans* infection. **(A)** Gene model of *RB* and splicing mutant *RB_IVS*. Red letters in *RB_IVS* denote mutations to change the intron splicing recognition sites. On the top is schematic of two transcripts of *RB* amplified using 5′-rapid amplification of cDNA ends (5′-RACE). UPM primer is Universal Primer A, GSP primer is a gene specific primer of *RB*. The white vertical lines represent single nucleotide polymorphisms that appeared in the sequencing results. The primers for *RB_CDS* transcript that cross exon-exon junctions are shown with dashed lines over the intron. Arrows denote the primers for alternatively spliced transcript *RB_IR* located on the intron sequence. The RT-PCR primers F and R are located on the two sides of the intron. The red arrows indicate the primers that are used to measure relative gene expression in Fig. S1B. The asterisk shows the location of a premature termination codon. On the right are diagrams showing the predicted proteins encoded by RB variants with domains indicated color. **(B)** RT-PCR analysis of *RB_IR* and *RB_CDS* in wildtype (WT), *RB* and *RB_IVS* transgenic *S. tuberosum* and *N. benthamiana*, PT29 (*S. bulbocastanum*); and AS events of *Rpi-sto1* in *Rpi-sto1* transgenic potato (*S. tuberosum*). *RB_IR* is 995 bp, and *RB_CDS* is 316 bp. *StEF1*α and *NbEF1*α were used as internal controls. **(C)** Expression of N terminal GFP-tagged fusions of RB, RB_CDS, and RB_IR in agroinfiltrated *N. benthamiana* was confirmed by protein blot using an anti-GFP antibody. The asterisks indicate the predicted sizes of RB_CDS and RB_IR. Equal protein loading is visualized by Ponceau S staining. **(D)** Splicing ratio of *RB* in transgenic potato following inoculation with water (mock), *P. infestans* JH19, or *P. capsici* LT1534. Transcript levels were detected by quantitative RT-PCR (RT-qPCR) and normalized to *RB_CDS* following mock treatment at 0 dpi. Bars represent the mean ± SD from three independent replicates. **(E)** RT-PCR of *RB_IR* or *RB_CDS* variants in *RB* transgenic potato following inoculation with different *P. infestans* strains (JH19, HB1501, T30-4), water (mock), or *P. capsici* strain LT1534. *StEF1*α was used as the internal control. The intensities under the PCR bands of each product were quantified by ImageJ software. **(F)** Splicing ratio of *RB* isoforms in *RB_IVS* transgenic potato following inoculation with *P. infestans* zoospores. RNA was extracted 0, 1, and 2 days post inoculation. Splicing ratio levels were detected by RT-qPCR. Bars represent the mean ± SD in three independent replicates. ns = no significant difference using a one-way ANOVA with Tukey’s test. **(G)** Splicing ratio of *RB* in *RB* transgenic *N. benthamiana* following agroinfiltration of RNAi-hairpin constructs then inoculation with *P. infestans* zoospores 2 days later. RNA was extracted at 0 and 2 days after pathogen inoculation. Splicing ratio levels were detected by RT-qPCR. Bars represent the mean ± SD in three independent replicates. **(H)** Splicing ratio of *RB* in *RB* transgenic *N. benthamiana* following treatment with DMSO (control), 1 µM PB, or 30 µM PB followed by inoculation with *P. infestans* zoospores 6 hours later. RNA was extracted at 0 and 2 days after pathogen inoculation. Splicing ratio levels were detected by RT-qPCR and bars represent the mean ± SD in three independent replicates. For graphs (D), (G), (H), ns, no significant difference, one asterisk denotes a significant difference at P<0.05, two asterisk denotes a significant difference at P<0.01 and three asterisks denotes a significant difference at P<0.001 using a two-way ANOVA with Tukey’s test. The above experiments were repeated 3 times with similar results.

To investigate the dynamic change of *RB* splicing during pathogen infection, we inoculated *RB* transgenic potato with *P. infestans* strain JH19 and measured the expression levels of different *RB* isoforms compared to levels following water (mock) inoculation. Quantitative RT-PCR data demonstrated that the splicing ratio (transcript level of the spliced isoform divided by the transcript level of the unspliced isoform) of *RB (RB_CDS/RB_IR)* significantly increased to 1.66 at 1 dpi and 2.78 at 2 dpi following *P. infestans* inoculation, indicating an increase in *RB_CDS* expression and a decrease of *RB_IR* expression during *P. infestans* infection (Fig. 1D and Supplemental Fig. S1B). Conversely, inoculation of *RB* transgenic potato with zoospores of *Phytophthora capsici* isolate LT1534 had no significant impact on *RB* splicing ratio alteration (Fig. 1D and Supplemental Fig. S1B). Following these experiments, the total gene expression of *RB* did not change significantly in either *P. infestans*-, *P. capsici*-, or mock-inoculated plants (Supplemental Fig. S1C). Previously, our results revealed that both the intron-spliced *RB_CDS* and intron-retained *RB_IR* can encode proteins when transiently expressed in *N. benthamiana* (Fig. 1C). A change in abundance of the two proteins was also observed during *P. infestans* infection. The abundance of the RB_CDS protein increased during *P. infestans* infection compared to mock treatment, whereas the accumulation of the truncated RB_IR decreased. However, no considerable change in protein expression level was noted during *P. capsici* infection (Supplemental Fig. S1D). These results indicate that the *RB* gene produces more RB_CDS due to increased intron splicing subsequent to *P. infestans* inoculation. To further verify the effect of *P. infestans* on *RB* AS, we tested multiple *P. infestans* genotypes. Genotype T30-4 and HB1501 had been sequenced and the presence of IPI-O1 had been reported (Thilliez et al., 2019; Zhang et al., 2021). Primers specific for IPI-O1 had been used for gene amplification in genotype JH19 and results in validation of functional IPI-O1. The RT-PCR results show that the relative expression intensity of the *RB_IR* isoform was reduced at 1 and 2 dpi, whereas the relative expression intensity of the *RB_CDS* isoform increased during the same timeframe, regardless of the pathogen genotype (Fig. 1E). The expression intensity remained unchanged in *P. capsici* inoculated plants and mock treatment (Fig. 1E). Collectively, these data supplemented that the *RB* gene produces more *RB_CDS* transcript due to increased intron splicing during *P. infestans* infection.

Given that *P. infestans* affects the expression level of *RB_CDS* via modulation of *RB* AS (Fig. 1D), we hypothesized that impeding the splicing process would impact *RB_CDS* expression. To test this, we assessed the *RB* splicing ratio change in splicing mutant *RB_IVS* transgenic potato during *P. infestans* infection. We found that the splicing ratio in this transgenic plant remained low and did not change significantly during infection, indicating that *RB* is maintained with the intron present (Fig. 1F and Supplemental Fig. S1E). In eukaryotes, the splicing process is executed by the spliceosome, a conserved ribonucleoprotein complex dynamically composed of various proteins (Chen and Manley, 2009). Among these components, U1 small nuclear ribonucleoprotein particle 70k (U1-70k) is crucial for binding the 5’splice site in pre-mRNA (Cho et al., 2011), and splicing factor 3b subunit 1(SF3b1) as the core component of the U2 snRNP, plays a pivotal role in branch site recognition (Will and Lührmann, 2011). Thus, we silenced these targets to verify the effect on splicing of *RB* transcripts. Additionally, as a control, we silenced the RNA-binding protein *GRP7*, known to regulate RNA processing, thereby affecting the PTI response and specifically influencing NLR protein Gpa2/Rx1-mediated immunity (Wang et al., 2020; Sukarta et al., 2022). The quantitative PCR results indicated that silencing of *U1-70k* and *SF3B1* both notably suppressed up-regulation of the *RB* splicing ratio following *P. infestans* infection compared to non-inoculated samples, whereas *GRP7* silencing did not affect the induction of *RB* splicing following infection (Fig. 1G and Supplemental Fig. S1, F and G). We also conducted a chemical splicing inhibitor assay using Pladienolide B (PB), a splicing modulator that targets SF3B1 and disrupts its proper recognition of the intron (Kotake et al., 2007; Cretu et al., 2018). The transcript expression analysis indicated that 30 μM PB effectively suppressed the splicing ratio increase during infection (Fig. 1H and Supplemental Fig. S1H). In conclusion, our results adequately demonstrated that the *RB_CDS* transcript was most prevalent following *P. infestans* infection, which is the consequence of an increase in intron splicing.

### Intron splicing of *RB* is necessary for the activation of resistance and functions independent of ETI

In light of the observed change in the *RB* splicing ratio during *P. infestans* infection, we aimed to characterize the function of different transcripts of *RB* in *RB*-mediated resistance. We transformed *RB_CDS*, *RB_IR*, and the splicing mutant *RB_IVS* into potato cultivar Favorita. The wild-type *RB* gene was also transformed into Favorita as a control. Three independent transgenic potato lines expressing intron-spliced transcript *RB_CDS* displayed resistance to *P. infestans*, similar to the three wild-type *RB* transgenic potato lines. In contrast, three independent lines expressing the intron-retained transcript *RB_IR* or three lines expressing splicing mutant *RB_IVS* all exhibited severe disease symptoms (Fig. 2A). The results of detached leaf assays of the transgenic potatoes were consistent with the whole plant phenotypes (Fig. 2, B and C). Additionally, we generated *RB*, *RB_CDS*, *RB_IR*, and *RB_IVS* transgenic *N. benthamiana* for our assays. Notably, only *RB_CDS* limited *P. infestans* lesion development (Supplemental Fig. S2, A to C). Furthermore, *RB_CDS*, neither *RB_IR* nor *RB_IVS*, induced HR in *N. benthamiana* in presence of its cognate effector IPI-O1 (Supplemental Fig. S2, D and E), further supporting that *RB_CDS* is required for *RB*-mediated immune responses. Of note, overexpression of RB_CDS and RB_IR did not induce HR in *N. benthamiana* (Supplemental Fig. S2, F to H).

**Figure 2.**
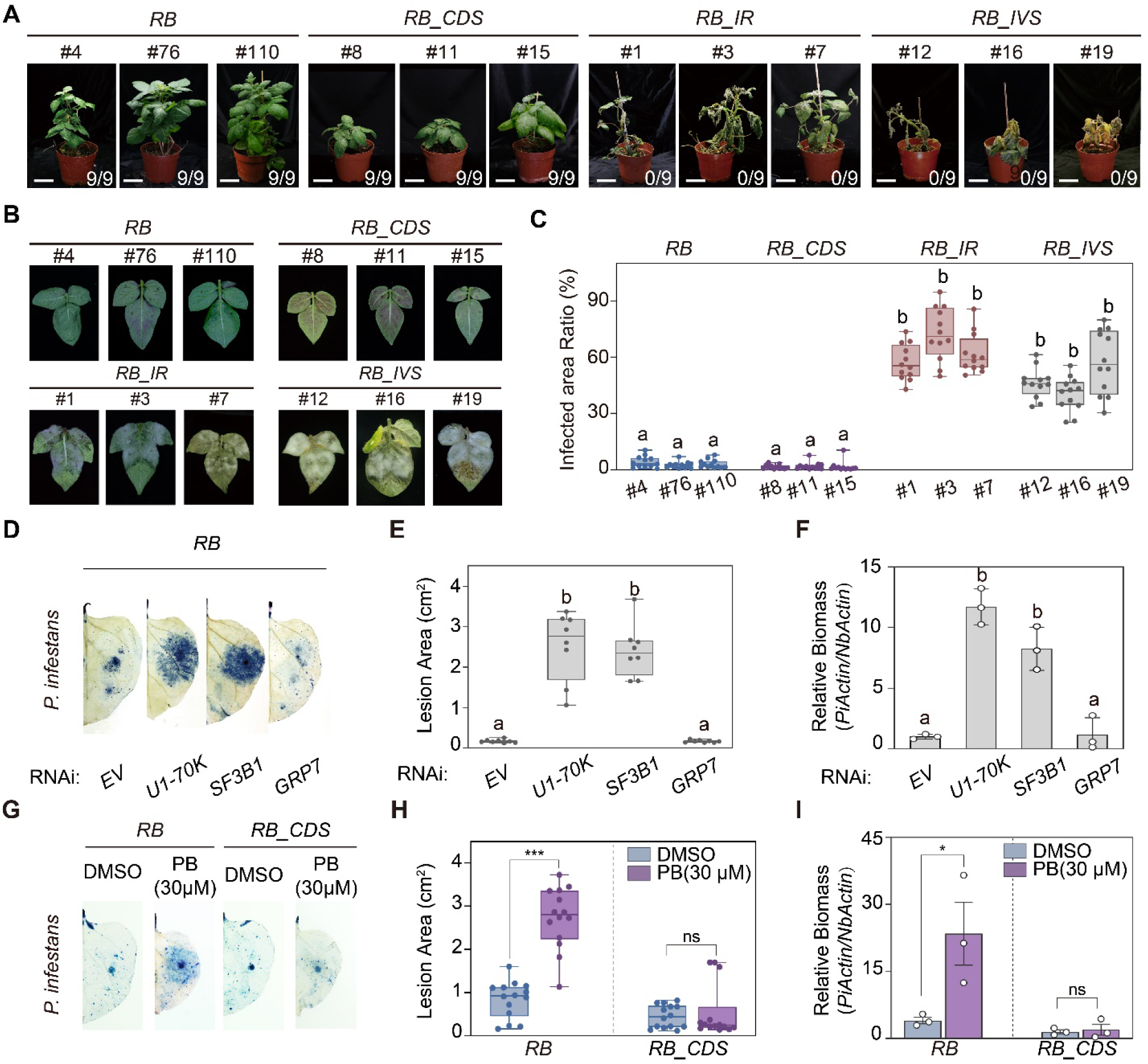
Intron retention in *RB* results in susceptibility to *P. infestans* infection. **(A)** A zoospore suspension of *P. infestans* strain JH19 was used to inoculate *RB*, *RB_CDS*, *RB_IR*, and *RB_IVS* transgenic potato plants at a concentration of 15,000 zoospores/mL. Plants were photographed at 5 dpi. For every transgenic potato, each line has nine repeats (n=9). The numbers represent the number of resistant plants compared to all inoculated plants. Bar=5 cm. **(B)** A zoospore suspension of *P. infestans* strain JH19 was drop-inoculated on detached leaves of *RB* transgenic potato expressing the indicated gene variants. The inoculated leaves were photographed at 4 dpi. **(C)** Infected areas (%) of inoculated leaves in (B). The infected area was measured in ImageJ. The data represent the mean ± SD from 12 replicates. Different letters denote a significant difference at P<0.001 using one-way ANOVA with Tukey’s HSD test. **(D)** Trypan blue staining of *RB* transgenic *N. benthamiana* following agroinfiltration of RNAi-hairpin constructs and inoculation with *P. infestans* zoospores. *EV* = empty vector control. Leaves were photographed at 5 dpi. (**E**) Lesion areas (cm^2^) of inoculated leaves in (D). Data represent the mean ± SD from 8 replicates. **(F)** Relative biomass of *P. infestans* was determined by quantitative PCR of *P. infestans* genomic DNA normalized to *N. benthamiana* genomic DNA at 5 days post inoculation (dpi). Bars represent the mean ± SD from three independent replicates. Different letters above bars denote a significant difference at P<0.001 using one-way ANOVA with Tukey’s HSD test. **(G)** Zoospores of *P. infestans* strain JH19 were inoculated on *RB* or *RB_CDS* transgenic *N. benthamiana* leaves pretreated with DMSO or 30 μM pladienolide B (PB). Leaves were stained with trypan blue and photographed at 5 dpi. **(H)** Lesion areas (cm^2^) of inoculated leaves in (D). Data represent the mean ± SD from 14 replicates. ns = no significant difference and three asterisks denotes a statistical difference at P<0.001 using a two-way ANOVA with Tukey’s test. **(I)** Relative biomass of *P. infestans* was determined by quantitative PCR of *P. infestans* genomic DNA normalized to *N. benthamiana* genomic DNA at 5 days post inoculation (dpi). Bars represent the mean ± SD from three independent replicates. ns = no significant difference, one asterisk denotes a significant difference at P<0.05 using a two-way ANOVA with Tukey’s test. The above experiments were repeated 3 times with similar results.

Given that only the protein encoded by the intron spliced transcript *RB_CDS* can activate immunity, we investigated whether interdiction of the splicing process might affect *RB*-mediated resistance to *P. infestans*. Silencing of *U1-70k, SF3B1*, and *GRP7* in *RB* transgenic *N. benthamiana* rendered *RB* transgenic *N. benthamiana* susceptible to *P. infestans* compared to the EV control, while silencing *GRP7* did not affect *RB*-mediated resistance (Fig. 2, D to F). Moreover, treatment with 30 μM PB significantly reduced *RB*-mediated resistance in *RB* transgenic *N. benthamiana*, but this treatment did not affect the resistance activation conferred by RB_CDS (Fig. 2, G to I). These results could adequately explain that the modulation of *RB* pre-mRNA intron splicing is essential for initiating *RB* resistance activation against *P. infestans*.

Most *R* genes function by recognizing their corresponding effectors and triggering a HR. To test whether disruption of *RB* splicing also interfered with elicitation of the HR, we co-expressed IPI-O1 and *RB* in *N. benthamiana* in which spliceosome components were silenced. Additionally, we introduced the intron-containing *R* gene *Rpi-blb2* (van der Vossen et al., 2005) and the intron-lacking *R* gene *R3a* (Huang et al., 2005) with their corresponding effector genes for HR testing. Following the silencing of *U1-70k* and *SF3B1*, co-infiltration of the intron-containing potato *NLR* genes *RB* or *Rpi-blb2* with their corresponding effectors *IPI-O1* and *Avrblb2*, respectively, resulted in a significantly weakened HR (Supplemental Fig. S3, A and B). Conversely, co-infiltration of the intronless *NLR* genes *RB_CDS* or *R3a* with the corresponding effectors *IPI-O1* and *Avr3a* induced a robust HR following the silencing of *U1-70k* and *SF3B1* (Supplemental Fig. S3, A and B). Silencing of *GRP7* had no effect on suppression the HR following co-infiltration of any tested NLRs with their corresponding effectors (Supplemental Fig. S3, A and B). Furthermore, treatment with 30 μM PB significantly reduced the incidence of HR upon IPI-O1 infiltration in *RB* transgenic *N. benthamiana*. However, this treatment did not affect the HR incidence conferred by RB_CDS (Supplemental Fig. S3C). These results suggest that the splicing plays a vital role in the activity of intron containing NLRs, including *RB* and *Rpi-blb2*.

### IPI-O1 mediates *RB* AS during the infection process

To determine the proteins secreted by *P. infestans* that regulate *RB* intron splicing, we designed an *RB* alternative splicing reporter system using the AS region of *RB* genomic sequences (from 1 nt to 1987 nt) fused with the *luciferase* (*LUC*) reporter gene (*35S::RB- LUC*) (Fig. 3A). This reporter system allows the detection of intron splicing modulation without ETI activation. The reporter system is predicted to produce two transcripts: the first is *RB_IR-LUC*, which retains the intron, introduces a premature termination codon, and does not produce a LUC signal; the second is *RB_CDS-LUC*, in which the intron is spliced and translates a complete RB CC-NB protein and functional LUC protein (Fig. 3A). The reporter gene β*-glucuronidase* (*GUS*) driven by *35S* promoter (*35S::GUS*) was used as an internal control (Fig. 3A). To verify intron splicing changes via the *RB* AS reporter system following *P. infestans* inoculation, we expressed *RB-LUC/GUS* via agrobacterium in *N. benthamiana* and subsequently inoculated with *P. infestans* JH19. Relative luciferase activity of *RB-LUC* was measured at 1 dpi and showed a significant increase following *P. infestans* treatment compared to the water-inoculated (mock). The relative luciferase activity of *RB_CDS-LUC* and *LUC* constructs remained unaltered following *P. infestans* inoculation (Supplemental Fig. S4). This verified the modulation of *RB* intron splicing by *P. infestans* using the *RB* AS reporter system, demonstrating the feasibility of this system. This supported the use of the *RB* AS reporter system in screening for *P. infestans* effectors that facilitate *RB* splicing. Indeed, in a previous study, we identified several splicing regulatory effectors (SRE) using another splicing reporter system (Huang et al., 2020). To assess the impact of SRE proteins on gene *RB* AS, we co-expressed several SRE proteins with the *RB* AS reporter system in *N. benthamiana*. Screening results show that IPI-O1 (SRE8) significantly induced *RB* intron splicing compared to the mock-infiltrated control (Fig. 3, B to D). Infiltration of SRE proteins with *RB_CDS-LUC*, which lacks the intron, did not affect LUC activity (Fig. 3, B to D).

**Figure 3.**
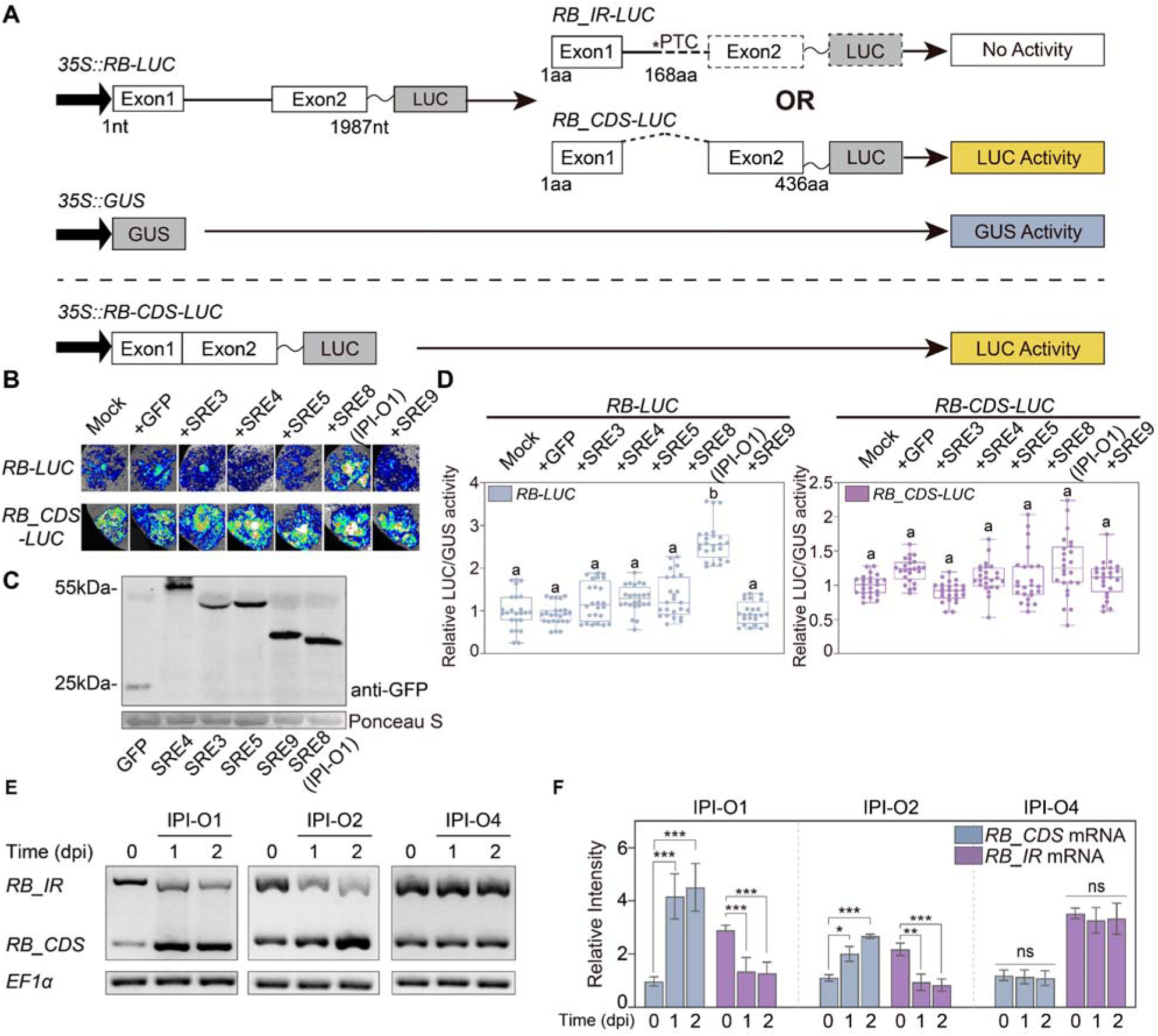
IPI-O1 induces *RB* intron splicing. **(A)** Schematic diagram of the *RB* alternative splicing reporter system. The genomic sequence of *RB* was fused with the *luciferase* (*LUC*) gene to construct *35S*::*RB-LUC*. Intron retention introduces a premature termination codon (PTC) and no production of the luciferase reporter. Intron splicing leads to an entire RB-LUC protein with luciferase activity. The *35S::GUS* construct was used as an internal control to normalize the activity of luciferase. The *35S*::*RB_CDS-LUC* construct with the intron removed produces a full-length RB-LUC protein with luciferase activity regardless of intron splicing activity. **(B)** Results of co-expression of the reporters with multiple splicing regulatory effectors (SRE) identified previously. *N. benthamiana* leaves were infiltrated with Agrobacterium carrying the indicated SRE protein. LUC activity was determined at 48 hpi using a chemiluminescent imaging system. **(C)** GFP-tagged SRE fusion protein expression in *N. benthamiana* was confirmed by protein blotting using an anti-GFP antibody. Equal protein loading was visualized by Ponceau S staining. **(D)** The LUC/GUS activity of infiltrated samples was calculated at 48 hpi. The relative LUC/GUS activities were relative to the activity of the construct with mock treatment. Data represent the mean ± SD from 24 replicates. Different letters above each bar represent a significant difference at P<0.001 using one-way ANOVA with Tukey’s HSD test. **(E)** Transcript levels of *RB_IR* and *RB_CDS* were determined by RT-PCR following agroinfiltration of IPI-O1, IPI-O2 and IPI-O4 into leaves of *RB* transgenic *N. benthamiana*. *NbEF1*α was used as the internal control. **(F)** Bars represent relative intensity of RT-PCR products in (D) and normalized with internal control *NbEF1*α. Intensities of the bands were quantified by ImageJ software, and the relative abundances of *RB* isoforms are relative to *RB_CDS* at 0 dpi. Bars represent the mean ± SD from three independent replicates. ns = no significant difference, one asterisk denotes a significant difference at P<0.05, two asterisks denotes a significant difference at P<0.01 and three asterisks denotes a significant difference at P<0.001 using a two-way ANOVA with Tukey’s test. The above experiments were repeated 3 times with similar results.

Classified within the *ipiO* locus in *P. infestans*, IPI-O1 is one of 16 variants in this family. Among them, IPI-O2 is highly similar to IPI-O1 and also induces HR with *RB*. The most divergent variant IPI-O4, differs from IPI-O1 by 20 amino acids and evades *RB* recognition (Champouret et al., 2009; Halterman et al., 2010). To test the ability of other IPI-O family members to modulate *RB* AS, we overexpressed these proteins in *RB* transgenic *N. benthamiana*. The followed RT-PCR of *RB* illustrated that, like IPI-O1, IPI-O2 induced *RB* intron splicing, but IPI-O4 does not (Fig. 3, E and F). Consistent with screening results by the *RB* AS reporter system, RT-PCR also revealed overexpression of other SRE proteins had no effect on *RB* splicing changes (Supplemental Fig. S5). Overall, it appears that IPI-O1 and its closely related family members can promote *RB* intron splicing, and this modulation correlates with *RB*-mediated immunity induction.

### Interaction between IPI-O1 and CWC15 is important for *RB* AS modulation

To identify the potential splicing regulators of *RB*, we performed an ULTImate (Hybrigenics) yeast-two hybrid (Y2H) screen to identify IPI-O1-interacting proteins within a *N. benthamiana* cDNA library. A total of 81.6 million interactions were analyzed and 254 clones were processed by sequencing of the prey inserts. Of these 254 clones, 28 contained an insert with homology to the *CWC15* gene from *N. tomentosiformis* (LOC104090615). Y2H assays have further verified the interaction of two full-length CWC15 homologs in *N. benthamiana* with IPI-O1 (Fig. 4A). Analysis of *CWC15* in potato revealed two *CWC15*-related gene sequences in the double-monoploid potato DM-1-3: *Soltu.DM.10G000900.1* (*StCWC15a*) and *Soltu.DM.07G027590.1* (*StCWC15b*), encoding 231 aa (amino acid) and 228 aa proteins respectively. StCWC15a and StCWC15b share 93% similarity at the amino acid levels (Supplemental Fig. S6A). Further analysis of CWC15 sequence variation in multiple wild and cultivated potatoes determined that CWC15 is well conserved (Supplemental Fig. S6B). CWC15 is a core component of the Nineteen complex within the spliceosome (Will and Lührmann, 2011), and is localized to the nucleus to modulate AS in Arabidopsis (Slane et al., 2020). To further test the interaction between IPI-O1 and StCWC15, we designed an IPI-O1 mutant based on a predicted NLS motif on the C terminal effector domain of IPI-O1 (Supplemental Fig. S7A). This mutant IPI-O1^M1^ contains a total of seven amino acids changes in a predicted NLS motif, where K (lysine) and R (arginine) were mutated to A (alanine) (Supplemental Fig. S7A). We performed co-immunoprecipitation in planta to test the interaction between IPI-O1 and mutant IPI-O1^M1^ with StCWC15a/b. The results showed that both StCWC15a and StCWC15b interacted with IPI-O1, but the mutant IPI-O1^M1^ failed to interact with either protein (Fig. 4B). As a control, we selected Pi04314, another RxLR effector that interacts with potato phosphatase PP1C and re-localizes the target from the nucleolus to the nucleoplasm (Boevink et al., 2016). No interaction was observed between StCWC15 and Pi04314. The direct interaction between StCWC15 and IPI-O1 was also verified *in vitro* by GST-pull down and Y2H. (Fig. 4C and Supplemental Fig. S7C).

**Figure 4.**
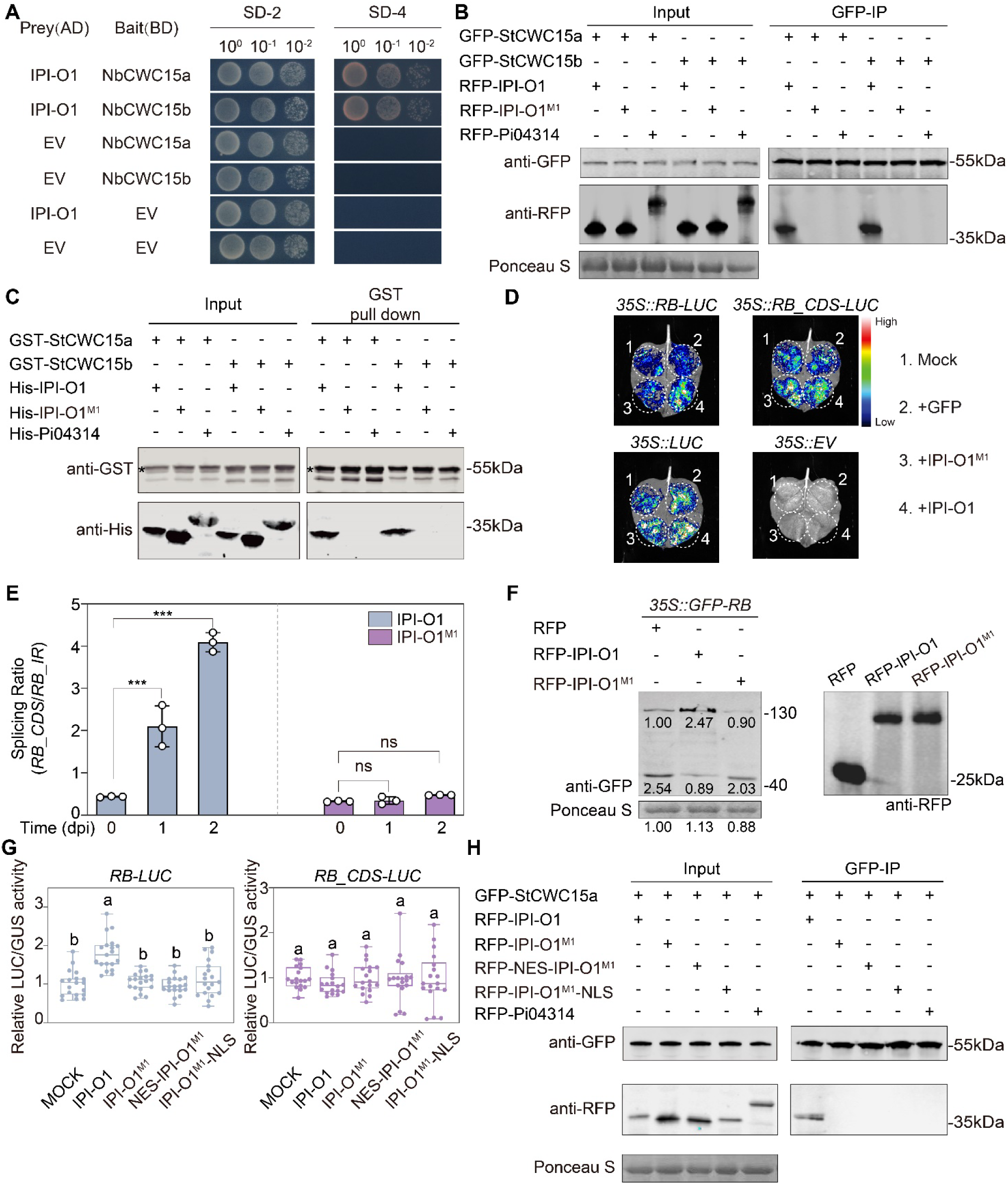
Interaction between IPI-O1 and StCWC15 is important for *RB* AS modulation. **(A)** Yeast-two hybrid interaction between IPI-O1 and CWC15 homologs from *N. benthamiana*. Yeast transformants were spotted as 10-fold serial dilutions on SD medium without leucine and tryptophan (SD–2), or without histidine, leucine, adenine, and tryptophan (SD–4). Yeast cells were photographed 2-3 days after plating. Empty BD and AD plasmids (EV) were used as controls. Experiments were performed at least three times with similar results. **(B)** Immunoprecipitation (IP) of StCWC15a/b-GFP with RFP-IPI-O1, RFP-IPI-O1^M1^, or RFP-Pi04314 following agroinfiltration in *N. benthamiana* leaves. Expression of constructs in the leaves is indicated by +. Markers are indicated on the right, and the Ponceau S staining was used to show levels of protein loading. **(C)** *In vitro* immunoprecipitation (IP) using the GST-tagged StCWC15a/b and IPI-O1 or IPI-O1^M1^. Input and GST-pull-down proteins were visualized by protein blotting using anti-GST and anti-His antibodies. Asterisks (*) indicate the protein bands that correspond to the predicted size of GST-CWC15a/b. **(D)** *N. benthamiana* leaves were infiltrated with Agrobacterium carrying the indicated constructs; relative LUC activity was measured at 48 hpi using a chemiluminescent imaging system. **(E)** Splicing ratio of *RB* in *RB* transgenic *N. benthamiana* following agroinfiltration of IPI-O1 and IPI-O1^M1^ at 0, 1 and 2 days by RT-qPCR. Bars represent the mean ± SD of three independent replicates. ns = no significant difference and three asterisks denote a statistical difference at P<0.001 using a two-way ANOVA with Tukey’s test. **(F)** The image on the left shows protein blot analysis of the protein accumulation of RB_CDS and RB_IR following transient expression of the indicated genes in *N. benthamiana* leaves. On the right, RFP tag fusions with IPI-O1 and IPI-O1^M1^ were confirmed by protein blotting using an anti-RFP antibody. Equal protein loading is visualized by Ponceau S staining. The intensities under the signal bands of each protein were quantified by ImageJ software. The above experiments were repeated 3 times with similar results. **(G)** The relative LUC/GUS activity of RB-LUC or RB_CDS-LUC with IPI-O1 related mutants was quantified at 2 dpi. The LUC/GUS activity of infiltrated samples was calculated by normalizing LUC activity to GUS activity. The LUC/GUS activities were relative to the activity of mock treatment. Data represent the mean ± SD from 19 replicates. Different letters above each bar represent a significant difference at P<0.001 using a one-way ANOVA with Tukey’s HSD test. **(H)** IPI-O1^M1^ fused NLS and NES motif cannot interact with StCWC15 in vivo by co-IP. Immunoprecipitation (IP) of StCWC15a-GFP with RFP-IPI-O1, RFP-IPI-O1^M1^, RFP-IPI-O1^M1^-NES and RFP-IPI-O1^M1^-NLS following agroinfiltration in *N. benthamiana* leaves. Expression of constructs in the leaves is indicated by +. Markers are indicated on the right, and the Ponceau S staining was used to show levels of protein loading.

To test the biological significance of StCWC15b and IPI-O1interaction, we expressed IPI-O1 and IPI-O1^M1^ in the *RB-LUC* reporter system to test AS modulation. The data showed that IPI-O1, but not IPI-O1^M1^, significantly increased the *RB* intron splicing (Fig. 4D and Supplemental Fig. S7D). Further evaluation of *RB* splicing ratio alterations using RT-qPCR demonstrated that IPI-O1 increased the *RB_CDS*/*RB_IR* splicing ratio, indicating elevated *RB_CDS* expression level and reduced *RB_IR* expression. Conversely, overexpression of IPI-O1^M1^ did not elicit a change in splicing ratio (Fig. 4E and Supplemental Fig. S7E). To detect the IPI-O1 and IPI-O1^M1^ modulation of RB protein expression levels, we co-expressed IPI-O1 and IPI-O1^M1^ separately with GFP-RB in *N. benthamiana* for immuno-blot analysis. The data demonstrated an increase in RB_CDS protein and a decrease in RB_IR protein only in the presence of IPI-O1, and not IPI-O1^M1^, when compared to GFP-RB expressed alone (Fig. 4F). These results confirmed that the IPI-O1 mutant IPI-O1^M1^ has lost the ability to modulate *RB* AS changes.

Using confocal microscopy, we found that mutant IPI-O1^M1^ failed to accumulate in the nucleoplasm and nucleolus, and localized primarily in the cytoplasm (Supplemental Fig. S7, F and G). which may also lead its inability to modulate *RB* AS. Thus, to rule out the possibility that this dysfunction is simply caused by a localization change, we fused a canonical NLS motif to the mutant IPI-O1^M1^, and fused a nuclear exporting signal (NES) as a control (Supplemental Fig. S7, F and G). The results show that overexpression of neither IPI-O1^M1^-NLS nor NES-IPI-O1^M1^ can promote *RB* intron splicing or increase the abundance of *RB_CDS* like IPI-O1^M1^ using *RB* AS reporter system (Fig. 4G). The results of interaction tests between NES-IPI-O1^M1^ and IPI-O1^M1^-NLS with StCWC15 indicated that neither mutant interacted with StCWC15 in vivo or in vitro (Fig. 4H and Supplemental Fig. S7H). Taken together, these results support that binding between CWC15 and IPI-O1 is important for its modulation of *RB* AS.

We have demonstrated that regulation of *RB* intron splicing is critical for resistance activation. To assess the ability of splicing mutant IPI-O1^M1^ to induce HR with RB, we expressed IPI-O1 in transgenic *N. benthamiana* expressing *RB* variants. The results showed that IPI-O1 induced the HR in both *RB* and *RB_CDS* transgenic *N. benthamiana*, but not in *RB_IR* plants (Supplemental Fig. S7, I and J). However, IPI-O1^M1^, IPI-O1^M1^-NLS, and NES-IPI-O1^M1^ were unable to induce the HR in *RB* transgenic *N. benthamiana*. Nevertheless, they were still able to elicit the HR in *RB_CDS* transgenic *N. benthamiana* (Supplemental Fig. S7, I and J). These findings collectively suggest that the interaction between IPI-O1 and splicing factor CWC15 is critical for modulating *RB* intron splicing, which is necessary for RB activation.

### IPI-O1 facilitates re-localization of StCWC15 into the nucleolus and speckles

To investigate the subcellular localization of CWC15 proteins in potato, we fused red fluorescent protein (RFP) to the C-terminus of StCWC15a and StCWC15b. While expression of fusion proteins in *N. benthamiana* demonstrated that StCWC15a primarily localized to the nucleoplasm (91%), the protein was occasionally found in the nucleolus (5%) and nuclear speckles (4%). StCWC15b was distributed similarly and was localized in the nucleoplasm (90%), nucleolus (5%), and nuclear speckles (5%). The IPI-O1 and StCWC15 homologs displayed similar localization expression alone (Fig. 5A). To investigate the subcellular localization of the interaction between IPI-O1 and StCWC15, we performed bimolecular fluorescence complementation (BiFC). The results demonstrated that IPI-O1 interacted with StCWC15 proteins in nuclear speckles and the nucleolus (Fig. 5B). We did not observe fluorescence when testing the interaction between StCWC15 proteins and IPI-O1^M1^ or Pi04314 (Fig. 5B). It is noteworthy that both the nucleolus and nuclear speckles are cellular structures highly correlated with RNA splicing (Liao and Regev, 2021). To investigate potential changes in StCWC15 subcellular localization influenced by IPI-O1, we co-expressed StCWC15a/b-RFP fusion proteins with GFP-IPI-O1 in *N. benthamiana*, with mutant GFP-IPI-O1^M1^ and GFP protein alone as controls. The results showed that StCWC15a protein was re-localized into speckles (63%) or the nucleolus (26%), and that StCWC15b protein was re-localized into speckles (60%) or the nucleolus (25%) when co-expressed with IPI-O1, but not with IPI-O1^M1^ or GFP (Fig. 5C and Supplemental Fig. S8A). To further investigate the re-localization of StCWC15 during *P. infestans* infection, *N. benthamiana* expressing GFP–CWC15a/b was inoculated with zoospores of transgenic *P. infestans* isolate 88069 expressing the fluorescent protein RFP. The number of cells with speckle-localized CWC15 near the infected hyphae increased more than in cells distant from the infected hyphae (Supplemental Fig. S8, B and C). This effect was not observed in cells expressing the GFP control during *P. infestans* infection (Supplemental Fig. S8, B and C). These findings suggest that IPI-O1 interacts with StCWC15 proteins in the nucleus and facilitates their re-localization from the nucleoplasm into speckles and the nucleolus.

**Figure 5.**
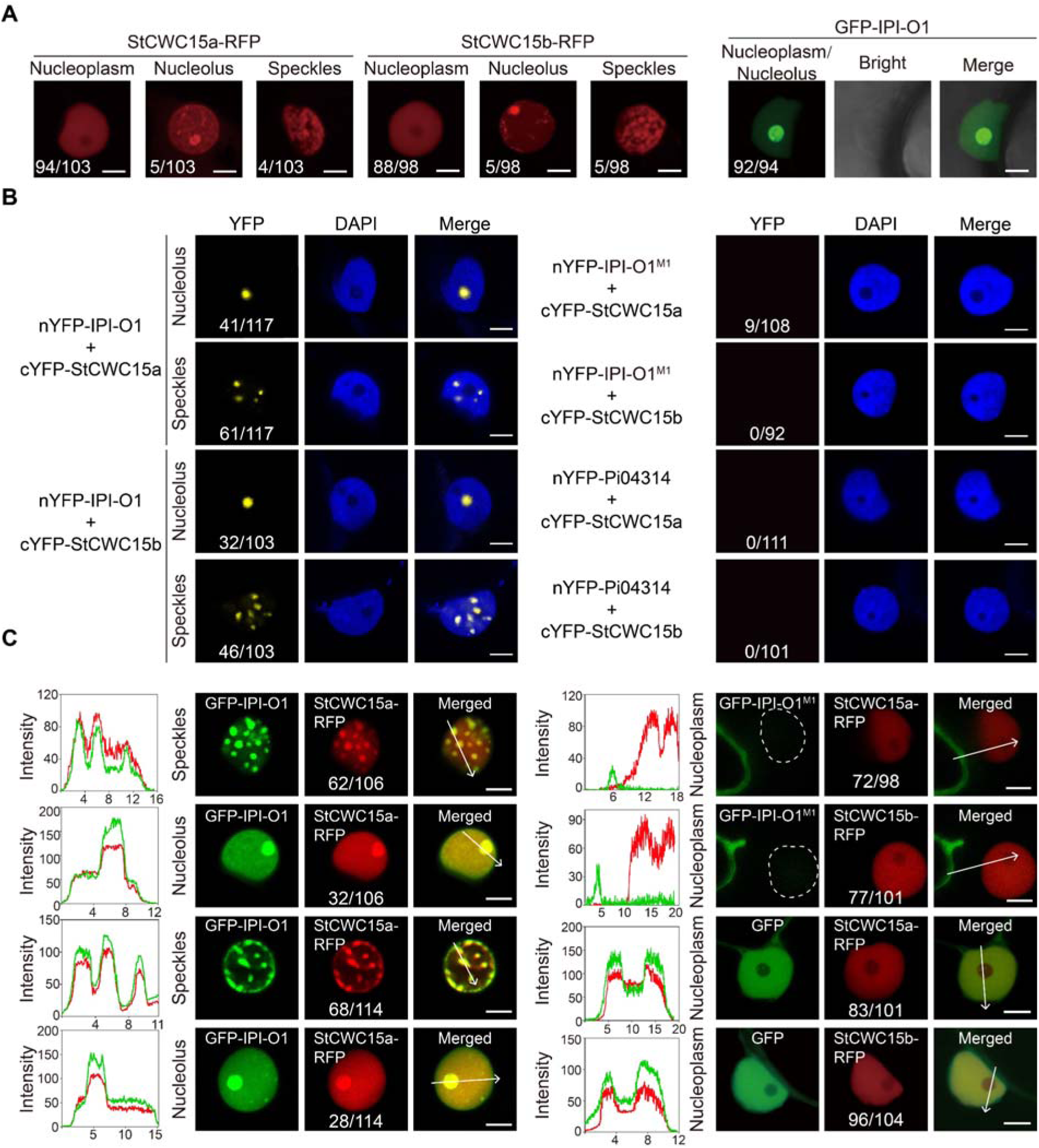
IPI-O1 re-localizes the StCWC15a/b from the nucleoplasm into speckles and nucleolus (A) StCWC15a/b-RFP or GFP-IPI-O1 were expressed in *N. benthamiana* leaves for 48 h before imaging. Numbers represent the frequency of subcellular localization in indicated cellular structure compared to the total fluorescent cells in the 25mm^2^ range. Bar = 5 μm. **(B)** IPI-O1 interacts with StCWC15 in the nucleolus and nuclear speckles. Split-YFP/BiFC of IPI-O1 or IPI-O1^M1^ and Pi04314 with StCWC15a and b using fluorescence imaging. Samples were stained with 2 µg /mL DAPI to indicate the nucleus. Numbers represent the frequency of cells showing YFP fluorescence compared to the total number of cells in the 25 mm^2^ discs from three biological replicates. Bar = 5 μm. **(C)** IPI-O1 co-localizes with StCWC15a/b in nuclear speckles and the nucleolus. GFP-IPI-O1, GFP-IPI-O1^M1^, and GFP alone were transiently co-expressed with StCWC15a/b-RFP by agroinfiltration in *N. benthamiana*. The left image is from the GFP channel, the middle image is from the RFP channel, and the right image is the overlay of the GFP and RFP channels. Numbers represent the frequency of cells showing indicated StCWC15 subcellular localization compared to the total StCWC15-positive cells in 25mm^2^ discs from three biological replicates. Graphs show the fluorescence intensity profiles across the arrows in the merged images. Bar = 5μm.

### CWC15 positively regulates *RB*-mediated HR and plant resistance against *P. infestans*

### infection

To determine the effect of CWC15 on *RB*-mediated immunity, we transiently silenced *CWC15*, transcription-related factor transcription initiation factor TFIID subunit 12b-like (*PPS7*) (Katou et al., 2005), splicing-related genes serine/Arginine-rich protein 30 (*SR30*), and arginine/serine-rich zinc knuckle-containing protein 32 (*RS2Z32*) (Reddy, 2004) in *RB* transgenic *N. benthamiana* followed by transient expression of IPI-O1. Infiltrated areas were assayed for HR elicitation and we found that silencing of *CWC15* resulted in a reduced incidence of HR compared to the EV control, while *PPS7*, *SR30*, or *RS2Z32* silencing had no effect on *RB* recognition of IPI-O1 (Supplemental Fig. S9A). *SGT1* silencing, known to be required for *RB* function (Bhaskar et al., 2008), also reduced HR incidence compared to the EV control (Supplemental Fig. S9A). The effect of CWC15 on *R* gene function appeared to be specific for *RB*, as silencing of *CWC15* had no effect on the elicitation of HR following co-infiltration of *R1*, *Rpiblb2*, or *R3a* with their corresponding effectors, or following infiltration of elicitor *INF* (Supplemental Fig. S9B).

Based on the transient screening results, we further verified the function of *CWC15* by virus-induced gene silencing (VIGS). The TRV-*NbCWC15* silenced plant exhibited small and short morphology compared to wildtype (Supplemental Fig. S10, A and B). Similar to the transiently silenced results, TRV-*NbCWC15* silenced plants disturbed the recognition of IPI-O1 by *RB* (Supplemental Fig. S10, C-E). Together, these indicated CWC15 as an interactor of IPI-O1 involved in *RB* activation. To verify the function of potato *CWC15* in *RB*-mediated resistance, we stably silenced *StCWC15* by introducing a hairpin-silencing construct into *RB*-transgenic potato by transformation. We obtained three independent transgenic lines in which *StCWC15* transcription was reduced and a susceptible phenotype was exhibited following inoculation with *P. infestans* when compared to the *RB* transgenic parent potato (Fig. 6, A to D). The morphology of *StCWC15* RNAi transgenic potato was similar to the observation in *N. benthamiana*, with a reduction in plant height and leaf size compared to *RB* transgenic potato (Supplemental Fig. S10F). CWC15 is able to regulate mRNA alternative splicing in *Arabidopsis* (Slane et al., 2020). To test whether StCWC15 could play a role in *P. infestans*-induced AS of *RB*, we detected *RB* transcript levels by RT-qPCR in the highest silencing efficiency line *RB-StCWC15* RNAi#2 following inoculation with *P. infestans* isolate JH19. We found that the *RB_CDS/RB_IR* splicing ratio remained constant in the *StCWC15* knockdown plant, while the ratio increased in the *RB* transgenic control plant (Fig. 6E). These data suggest that CWC15 can modulate *RB* splicing, thereby influencing *RB*-mediated immunity.

**Figure 6.**
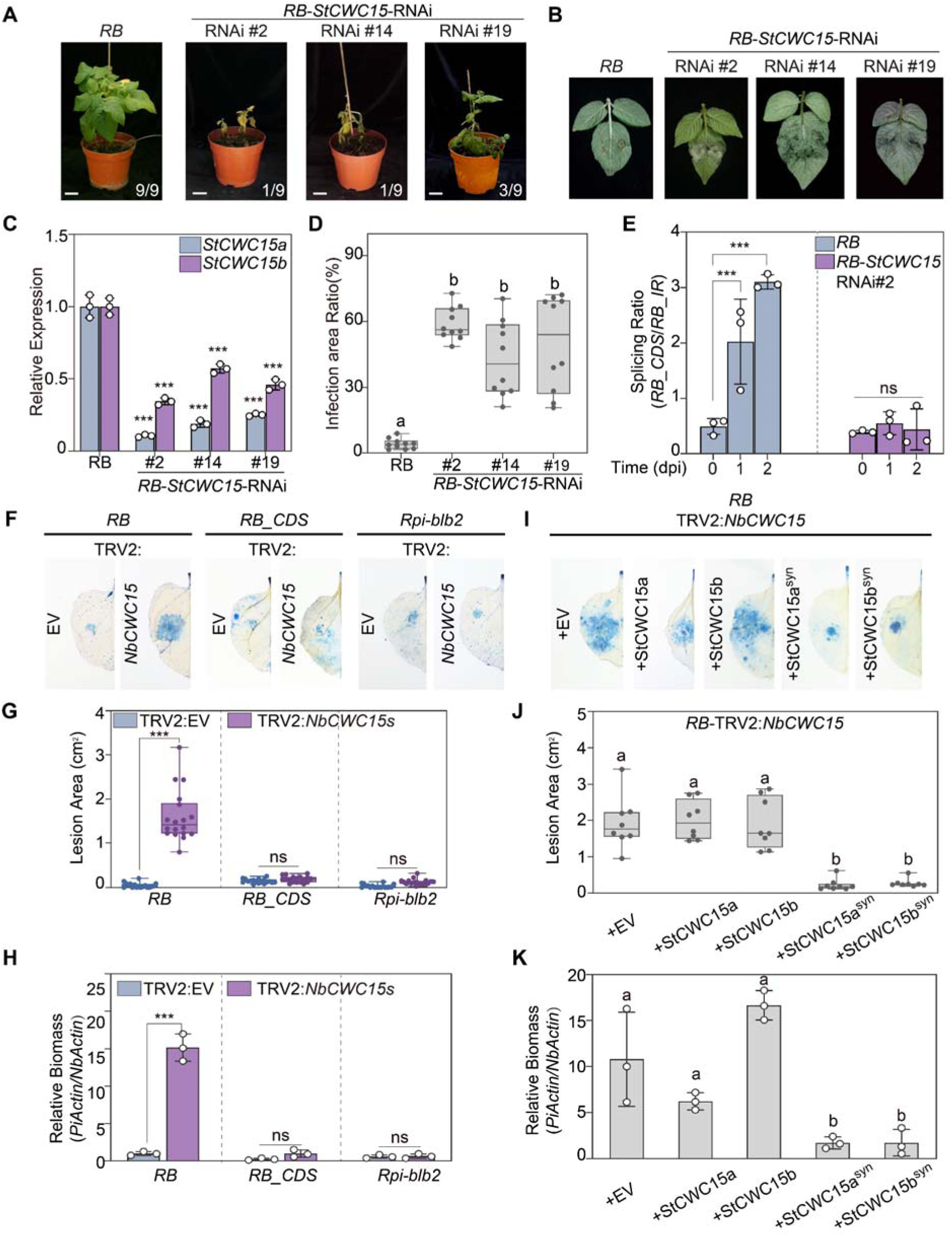
Silencing *StCWC15* disrupts *RB*-mediated resistance. **(A)** Whole plants of *RB* and *RB-StCWC15* RNAi transgenic were spray-inoculated with zoospores of *P. infestans* strain JH19 at a concentration of 15,000 zoospores/mL. Plants were photographed at 5 dpi. Inoculation of each transgenic potato line was repeated nine times (n=9). Bar = 3 cm. The numbers represent resistant plants compared to all inoculated plants. **(B)** Zoospore suspension of *P. infestans* strain JH19 was drop-inoculated on detached leaves of *RB* and *RB-StCWC15* RNAi transgenic potato lines. The infected leaves were photographed at 4 dpi. The numbers represent the number of resistant leaves compared to all inoculated leaves. **(C)** Relative expression of *StCWC15a/b* in three RNAi transformed lines compared to *RB* transgenic potato. Bars represent the mean ± SD of three independent replicates. Three asterisks designate a statistical difference from the *RB* control at P<0.001 using a two-way ANOVA with Tukey’s test. **(D)** Infected areas (%) of inoculated leaves in (B). The infected area was measured in ImageJ. The data represent the mean ± SD from 10 replicates. Different letters denote a significant difference at P<0.001 using a one-way ANOVA with Tukey’s HSD test. **(E)** Splicing ratio of *RB* in *RB-StCWC15* RNAi transgenic potato line#2 following inoculation with *P. infestans* zoospores at 2 dpi by RT-qPCR. Bars represent the mean ± SD from three independent replicates. ns=no significant difference and three asterisks denote a statistical difference at P<0.001 using a two-way ANOVA with Tukey’s test. **(F)** *NbCWC15* was silenced by VIGS in *RB*, *RB_CDS*, and *Rpi_blb2* transgenic *N. benthamiana* and inoculated with *P. infestans* JH19 on detached leaves. Leaves were stained with trypan blue and photographed at 5 dpi. **(G)** Lesion areas(cm^2^) of inoculated leaves in (F) were calculated using ImageJ. Data represent the mean ± SD from 18 replicates. **(H)** Relative biomass of *P. infestans* was determined by qPCR of *P. infestans* genomic DNA normalized to *N. benthamiana* genomic DNA at 5 days post inoculation (dpi). Bars represent the mean ± SD from three independent replicates. Three asterisks designate a statistical difference from the control at P<0.001 using a two-way ANOVA with Tukey’s test. **(I)** Complementation of *RB* function caused by *CWC15* silencing in *N. benthamiana* via synthetic StCWC15 constructs (StCWC15a/b^syn^). Agrobacterium expressing StCWC15a/b^syn^ constructs were infiltrated into *NbCWC15* silenced *RB* transgenic *N. benthamiana*, then inoculated with *P. infestans* JH19 zoospores. Leaves were stained with trypan blue and photographed at 5 dpi. **(J)** Lesion areas(cm^2^) of inoculated leaves in (I) were calculated using ImageJ. Data represent the mean ± SD from 8 replicates. **(K)** Relative biomass of *P. infestans* was determined by qPCR of *P. infestans* genomic DNA normalized to *N. benthamiana* genomic DNA at 5 days post inoculation (dpi). Bars represent the mean± SD from three independent replicates. Different letters denote a significance difference at P<0.001 using one-way ANOVA with Tukey’s HSD test.

To detect the possibility of CWC15 involvement in the recognition of IPI-O1 by RB_CDS as a tradition guardee protein. We assayed for *P. infestans* resistance and HR induction following silencing *CWC15* in *RB*, *RB_CDS* and *Rpi-blb2* transgenic *N. benthamiana*. Silencing of *CWC15* significantly impaired IPI-O1 induced HR and resistance to *P. infestans* in the *RB* line, but had no effect on the resistance in *RB_CDS* and *Rpi-blb2* transgenic lines (Fig. 6, F to H). Additionally, *CWC15* silencing had no effect on the HR induced by RB_CDS/IPI-O1, Avr3a/R3a or INF1 (Supplemental Fig. S11, A and B). This effect does not appear to be due to interaction between CWC15 and RB, as no interaction was found with RB_CDS or RB_IR proteins (Supplemental Fig. S11C). These data highlight that splicing factor CWC15 is specifically involved in regulating *RB* splicing changes during *P. infestans* infection, effecting *RB*-mediated immunity in potatoes, but does not affect *RB* function once the intron is removed. In further experiments, we complemented *StCWC15* in the TRV-*NbCWC15* silenced *RB* transgenic *N. benthamiana* using silencing-resistant *CWC15* synthetic constructs to test CWC15 function. These synthetic genes were constructed with shuffled synonymous codons and encode full-length StCWC15a/b (Supplemental Fig. S12A). Expression of StCWC15a^syn^ and StCWC15b^syn^ rescued the impaired HR and *RB* resistance in TRV silenced-*CWC15* transgenic plants (Fig. 6, I to K; Supplemental Fig. S12B). Immunoblot analysis revealed that expression of StCWC15a^syn^ and StCWC15b^syn^ recovered the expression of CWC15 in silenced *N. benthamiana* (Supplemental Fig. S12, C and D). Collectively, our results indicate that IPI-O1 interacts with the splicing factor CWC15 to mediate the AS of *RB*, leading to increased expression of resistant transcript *RB_CDS* and activation of *RB*-mediated immunity.

### Regulation of *RB* intron splicing is critical for balancing plant growth and immunity

Our prior findings indicated that the *RB* intron is primarily unspliced in the absence of *P. infestans*, leading to the production of mainly truncated protein. The presence of *P. infestans* effector IPI-O1 in the nucleus leads to increased intron splicing and increased translation of RB_CDS protein. This discovery suggests that the plant maintains resistance gene *RB* in a non-spliced state during normal growth and may avoid overexpression RB_CDS inappropriate activation of immunity responses that could impact growth and development. To test this, we generated *RB*, *RB_CDS*, and *RB_IR* transgenic potatoes with a CaMV 35S promoter to assay their growth characteristics. Firstly, we screened the positive transgenic lines that successfully inserted the target gene from multiple differentiated clustered buds using PCR verification. The expression level of *RB_CDS* differs in multiple *35S::RB_CDS* transgenic lines compared to the *RB* transgene driven by the native promoter (Supplemental Fig. S13A). Three lines with a single-copy of the transgene (#8, #11, #15) were identified among 34 *RB_CDS* transformants and showed a 15-20 fold increase of *RB_CDS* expression compared to the native promoter *RB* (Supplemental Fig. S13A). To ensure consistency of gene copy number and expression level in the various *RB* transgenic potatoes, we identified three single-copy transgene lines each from 67 *RB*-transformed seedlings and 29 *RB_IR*-transformed seedlings with similar gene transcription levels (Supplemental Fig. S13B and Supplemental Data Set S5). *RB_CDS* transgenic lines exhibited shorter stem height and root length than non-transformed Favorita in the seedling stage, while *RB* and *RB_IR* transgenic lines were not significantly different (Fig. 7, A-C). Furthermore, *RB_CDS* transgenic potato also displayed slower growth and dwarf phenotypes compared to the wild-type *RB* transgenic potatoes when grown in pots (Supplemental Fig. S13, C and D). The total tuber quality per plant and the individual tuber quality were significantly reduced in plants overexpressing *RB_CDS* compared with wildtype, while total tuber quality and individual tuber quality in *RB* and *RB_IR* transgenic potatoes were not significantly altered (Fig. 7, D and E, Supplemental Fig. S13E). Studies have shown that accumulation of salicylic acid (SA) in response to biotic stress affects the growth of plants (Ning et al., 2017; Gao et al., 2021a; Gao et al., 2021b). We compared SA concentrations in *RB* and *RB_CDS* transgenic potatoes with non-transformed wild-type potatoes and found that SA accumulation in the *RB_CDS* transgenic line was higher than in wild-type and *RB* transgenic potatoes (Fig. 7F). Additionally, the transcription of *SAR Deficient 1* (*SARD1*), a key regulator of SA synthesis and plant immunity (Zhang et al., 2010), and the SA-regulated gene *pathogenesis-related gene 1* (*PR1*) were up-regulated in *RB_CDS* transgenic potatoes compared to wild-type Favorita and *RB* transgenic potatoes (Fig. 7G and H). These results indicate that the potatoes sustained overexpression of *RB_CDS* leads to continuous activation of SA-mediated defense responses, which likely affects plant growth and tuber production.

**Figure 7.**
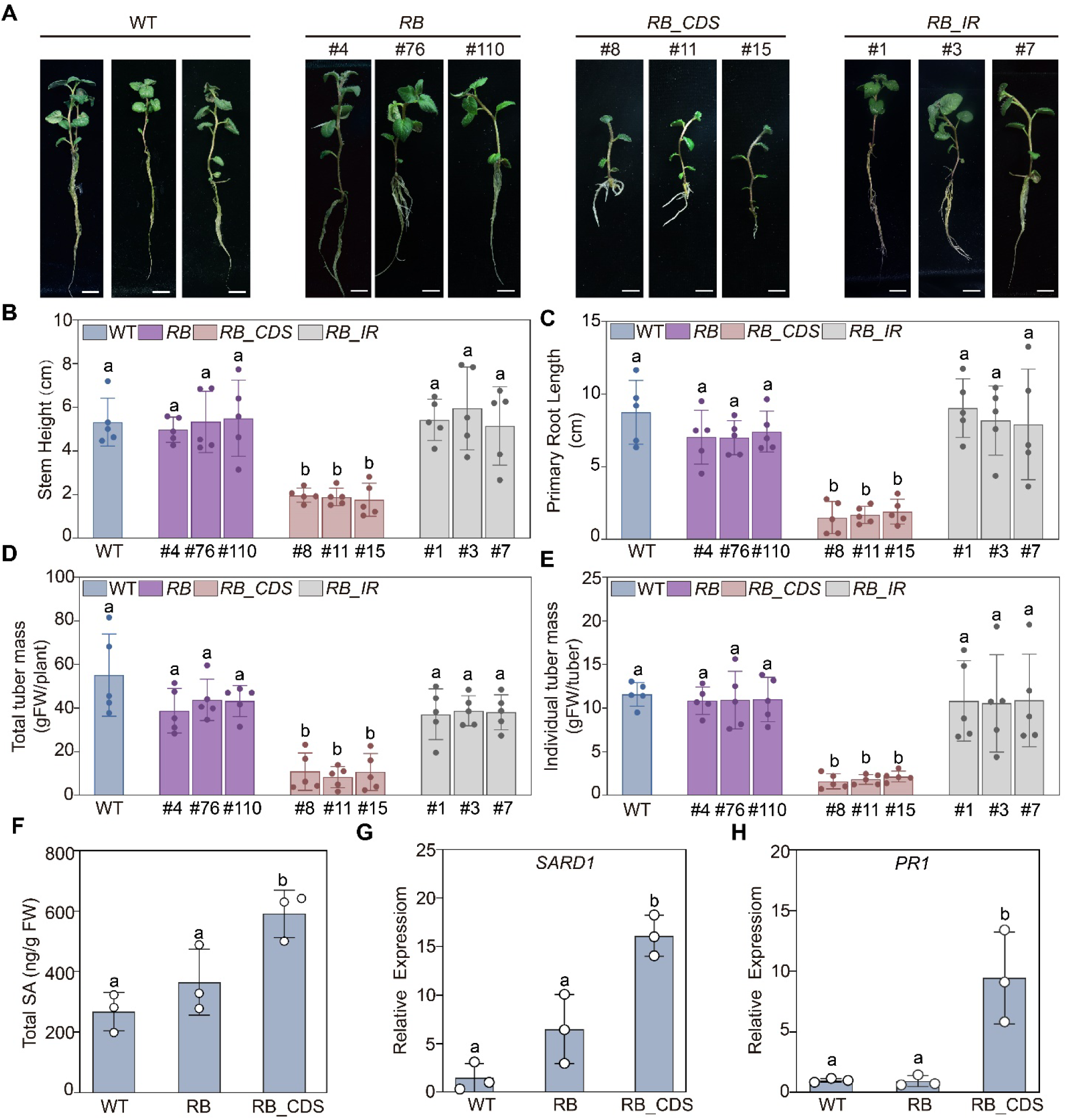
Alternative intron splicing of *RB* balances plant growth and immunity. **(A)** Photos of 21-day-old wild-type (WT) Favorita and *RB*, *RB_CDS*, and *RB_IR* transgenic tissue culture plants. Bar = 1 cm. **(B)** Average stem height and **(C)** root length of Favorita (WT) and 21-day-old transgenic potato tissue culture plants. Bars represent the mean ± SD from 5 tissue culture plants of 3 independent lines for each genotype. Different letters represent a statistical difference at P<0.01 using one-way ANOVA with Tukey’s HSD test. **(D)** Total tuber mass and **(E)** individual tuber mass of tubers from greenhouse grown plants. Measurements were taken at 8 to 10 weeks after tissue culture plants were transplanted. Bars represent the mean ± SD from 5 replicates of 3 independent lines for each genotype. Different letters represent a statistical difference at P<0.01 using one-way ANOVA with Tukey’s HSD test. **(F)** Total salicylic acid (SA) content in Favorita (WT) and transgenic *RB* and *RB_CDS* lines without pathogen inoculation. FW, fresh weight. Bars represent the mean ± SD from three independent replicates. Different letters represent a statistical difference at P<0.05 using one-way ANOVA with Tukey’s HSD test. **(G-H)** Relative expression of *SARD1* (G) and *PR1* (H) in the indicated transgenic potato as determined by RT-qPCR. Bars represent the mean ± SD from three independent replicates. Different letters represent a statistical difference at P<0.01 using one-way ANOVA with Tukey’s HSD test.

Previous reports have shown that truncated NLR proteins are expressed and have a potential role in plant immunity (Nishimura et al., 2017; Liang et al., 2019). Especially the C-terminally truncated protein, which are encoded CC or TIR of NLR, their self-association is critical for their activation of downstream immunity (Rairdan et al., 2008; Gutierrez et al., 2010; Maekawa et al., 2011). In this work, we identified RB encoded a truncated NLR protein RB_IR during normal growth stage, which encompassed the N-terminal of RB but cannot activate immunity (Fig. 2A). To further explore the interplay of truncated and full-length RB proteins, we tested the effect of RB_IR on RB_CDS-mediated immunity. We overexpressed *RB_IR* in *RB_CDS* transgenic *N. benthamiana* followed by inoculation with *P. infestans*. The results show that overexpression of RB_IR compromised RB_CDS resistance compared to the GFP control, whereas overexpression of R2-CC, which encodes the N-terminus of R2, had no effect on RB_CDS resistance (Fig.8, A to C). Overexpression of RB_IR also disrupted the HR induced by IPI-O1 in *RB_CDS* transgenic *N. benthamiana* (Supplemental Fig. S14), indicating that RB_IR is likely to disturb the RB-mediated resistance. The N terminal CC domain of RB displayed self-interaction *in vivo* and *in vitro*, and is vital for immunity activation (Chen et al., 2012; Zhao and Song, 2021). To detect the interaction relationship between RB_IR and RB_CDS, we transiently co-expressed them in *N. benthamiana* and performed co-IP The result showed that RB_IR can interact with full-length RB_CDS, as well as the N terminal CC domain of RB (Fig. 8D). Together, these results suggest that truncated protein RB_IR can interact with the CC domain of RB_CDS, and that its overexpression affects RB_CDS mediated resistance. This is likely due to the disrupt of RB-CC self-association caused by an elevated amount of RB_IR, as we observed that overexpression of RB_IR could affect the self-association of RB-CC through competitive binding to RB_CC protein (Fig. 8E). The same concentration of another N-terminal protein, R2-CC, had no effect on either RB_CDS immunity activation or RB-CC self-association (Fig. 8E). Together, these findings suggest that the overexpression of RB_IR could potentially compete with N-terminal self-association of full-length RB protein, consequently disrupting immunity activation.

**Figure 8.**
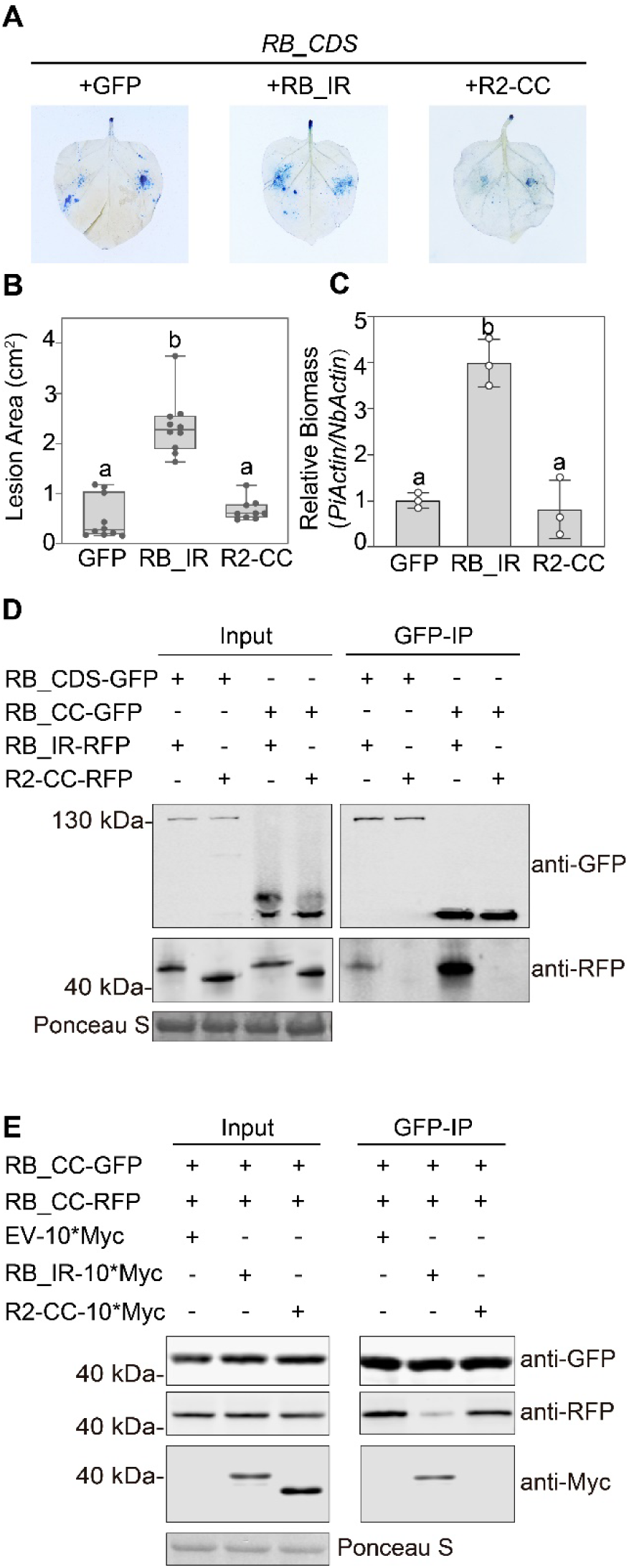
Overexpression of RB_IR disrupts RB_CDS N-terminal self-association and immunity activation. **(A)** Zoospores of *P. infestans* strain JH19 were inoculated on *RB_CDS* transgenic *N. benthamiana* leaves expressing the indicated proteins by agroinfiltration. Leaves were stained with trypan blue and photographed at 5 dpi. **(B)** Lesion areas (cm^2^) of inoculated leaves in (B) were calculated using ImageJ. Data represent the mean ± SD from 10 replicates from three independent experiments. **(C)** Relative biomass of *P. infestans* was determined by qPCR of *P. infestans* genomic DNA normalized to *N. benthamiana* genomic DNA at 5 dpi. Bars represent the mean ± SD from three independent replicates. For B and C graphs, different letters represent a statistical difference at P<0.001 using one-way ANOVA with Tukey’s HSD test. **(D)** Immunoprecipitation (IP) of RB_CDS-GFP and RB-CC-GFP with RB_IR-RFP, R2-CC-RFP following agroinfiltration in *N. benthamiana* leaves. Expression of constructs in the leaves is indicated by + Ponceau S staining was used to show levels of protein loading**. (E)** Agrobacterium carrying RB-CC-GFP and RB-CC-RFP (OD_600_ = 0.3) were co-expressed in *N. benthamiana* in the presence of EV-10×Myc empty vector (EV-10*Myc represents pBIN empty vector fused with 10×Myc tag), RB_IR-10×Myc, or R2-CC-10×Myc (OD_600_ = 0.6) by agroinfiltration. Total protein was incubated with GFP-Trap beads for co-IP assays, and immunoprecipitated proteins were analyzed using anti-GFP, anti-RFP, or anti-Myc antibodies. Ponceau S staining was used to show levels of protein loading. These experiments were repeated three times with similar results.

## Discussion

Earlier studies indicate interaction between IPI-O1 and the RB CC domain in vivo (Chen et al., 2012). Further evidence indicates that IPI-O1 accumulates in the nucleus and it induces the HR with RB protein in the cytoplasm (Wang et al., 2018; Zhao and Song, 2021). Our current work builds on previous data and uncovers a new model of NLR mediated plant immunity responses: a pathogen effector (IPI-O1) targets a component of the spliceosome (CWC15) in the nucleus, modulating the intron splicing of a plant NLR gene (*RB*). This results in increased expression of the functional transcript *RB_CDS* that ultimately leads to resistance activation. In addition, this study implicates the spliceosome as a regulatory complex in the plant that can surveil pathogen effector activity and activate plant immunity through proper splicing of an NLR. Meanwhile, alternative intron splicing of the *NLR* regulates expression to prevent inappropriate stress-induced responses during normal growth while maintaining the ability to rapidly activate plant immunity upon detection of the pathogen (Fig. 9).

**Fig. 9.**
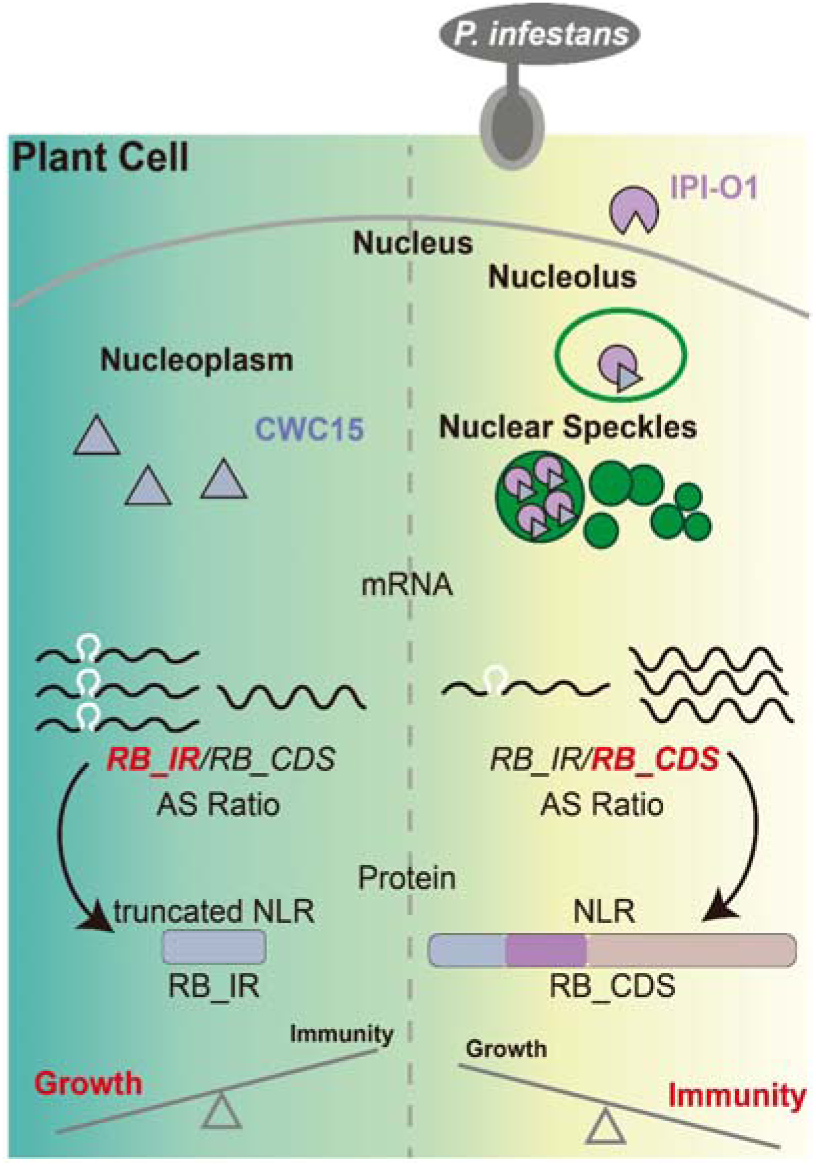
Schematic model of the P. infestans host pathosystem related to resistance gene RB and effector IPI-O1. During normal growth (left), CWC15 proteins (grey triangles) are maintained in the nucleoplasm and *RB* transcripts exist primarily with the intron retained (*RB_IR*). This results in the production of a truncated, non-functional RB protein that supports growth over immunity. Following *P. infestans* infection, effector IPI-O1 is introduced into the plant cell and translocated to the nucleus, leading to a re-localization of CWC15 from the nucleoplasm to the nucleolus and speckles. This results in increased splicing of the *RB* intron (*RB_CDS*), resulting in production of a full-length RB protein that can elicit immunity.

Pathogens and plants are eternally entrenched in molecular warfare, utilizing an arsenal of microbial effector proteins that manipulate the host to cause disease and an equally diverse set of host protein receptors that recognize the presence of pathogen effectors to activate immune responses. Given the complexity of possible interactions between host and pathogen molecules, it is no surprise that multiple mechanisms exist to mediate effector recognition and activate resistance. Classical activation models of NLR proteins and effectors include direct recognition, indirect recognition, integrated decoy, and activation traps (Duxbury et al., 2021). The simplest mechanism is direct interaction between NLR proteins and their corresponding effector, leading to changes in protein structure or location that results in resistance activation (Dodds et al., 2006). The indirect recognition mechanisms are divided into the guardee and the decoy models. In the guardee model, the effectors target a defense-related protein, either through direct binding or protein modification, that is monitored by an NLR protein. When changes in the guarded protein are recognized, this leads to activation of the NLR protein and elicitation of immune responses (Dangl and Jones, 2001; Axtell and Staskawicz, 2003). The decoy model is derived from the guardee model, but decoy proteins have no function in host defense or susceptibility in the absence of the NLR protein (van der Hoorn and Kamoun, 2008; Zhou and Chai, 2008). Our results demonstrate an additional mechanism of effector function to modulate *NLR* expression in plants. The proposed model involves the spliceosome component CWC15 protein, which is translocated to the nucleolus and nuclear speckles in the presence of IPI-O1, leading to splicing of the intron within the *RB* gene. The outcome of this model is similar to TAL effector activation traps that result in increased transcription of multiple executor *R* genes (Bogdanove et al., 2010). However, in the case of IPI-O1, we found that total *RB* transcription is not significantly induced during infection. This is inconsistent with the results of previous studies that demonstrated an increase in *RB* transcription during *P. infestans* infection in potato (Kramer et al., 2009), but is consistent with reports that *RB* is transcribed at a constitutive level (Bradeen et al., 2009). These contrasting observations are most likely due to differences in host and pathogen isolates with different genotypes used in our experiments, which could affect IPI-O1 expression and *RB* transcript accumulation. The primers and methods used for detection of RB transcripts was also different between these studies. Regardless, our results indicate that *RB* gene expression is likely regulated at the co-transcriptional or post-transcriptional level, and dose not conform to a transcription activation model through promoter binding as reported in other pathosystems such as with TAL effectors.

IPI-O1 is one of multiple *P. infestans* effectors capable of reprogramming genome-wide alternative intron splicing of the host (Wang et al., 2018; Huang et al., 2020; Wang et al., 2023). Consistent with previous results, we observed that IPI-O1 was localized in the nucleus and further highlighted the function of IPI-O1 in the nucleus. We clearly revealed that IPI-O1 can interact with the splicing factor CWC15 in speckles and the nucleolus, which are known to be sites for pre-mRNA splicing processing (Han et al., 2011; Liao and Regev, 2021). The interaction is important for IPI-O1 modulation of *RB* AS, which is crucial for activating resistance. Although we have focused on AS of *RB*, it seems unlikely that *RB* is the specific target of IPI-O1. More likely, IPI-O1 interaction with the spliceosome affects AS at a global level. This is already supported by previous data showing that IPI-O1 regulate AS of immunity-related gene *RLPK* (Huang et al., 2020). The impact of IPI-O1/CWC15 complex on other genes AS regulation will be a focus of future research. The other study showed that IPI-O1 is localized to both the cytoplasm and the nucleus, this discrepancy in localization of IPI-O1 may be due to GFP-tag fusion on the different positions of IPI-O1, which could affect the protein structure. In Zhao et al.’s work, the cytoplasmic localization of IPI-O1, was found to be important for recognition by the RB (Zhao and Song, 2021). We believe that, like many effectors, both subcellular localizations in the nucleus and cytoplasm are necessary for resistance activation (Lüdke et al., 2022).

NLR proteins are the intracellular immune receptors and activators in plants. The transition between autoinhibited and activated states of NLRs is fine-tuned by intra- and inter-molecular interactions. Over-accumulation of R proteins could lead to autoimmunity (Dinh et al., 2022), whereas insufficient NLR expression could lead to pathogen susceptibility. The expression of *NLR* genes is regulated at different levels, including transcriptional level, post-transcriptional level and protein level (Li et al., 2015). AS serves as a key player in gene regulation at the RNA level and has been recognized for its role in regulating plant immunity (Reddy, 2007; Huang et al., 2017; Reddy et al., 2020). Apart from global role of AS in regulating plant disease resistance responses, it also directly affecting the expression of *NLR* genes. *Arabidopsis snc1* mutants constitutively activate defense responses and have increased resistance to pathogens (Zhang and Li, 2005). A screen for suppressors of *snc1* revealed two *MOS* (modifier of *snc1*) genes, *MOS4* and *MOS14*, that are involved in mRNA splicing (Palma et al., 2007; Xu et al., 2011). The MOS14 protein is similar to transportin-SR, which is necessary for nuclear import of serine-arginine (SR) rich proteins. The *mos14-1* in *Arabidopsis* affects splicing of *NLR* genes *SNC1* and *RPS4* and compromises RPS4-dependent immunity and basal resistance to *Pseudomonas syringae* DC3000 (Xu et al., 2011). In *Arabidopsis*, *P. syringae* introduces small RNAs that silence *SWP73A*, a chromatin-remodeling gene that regulates expression of several *NLRs* through promoter binding and alternative splicing (Huang et al., 2021). *CDC5* has been reported to regulate alternative splicing of *RPS4*, which is important for RPS4 activity (Palma et al., 2007; Zhang and Gassmann, 2007). In this work, the IPI-O1 effector interacts with CWC15 to regulate splicing of the *RB* gene. It seems unlikely that *RB* is only regulated by IPI-O1. More likely, the CWC15 and RB intron can be regulated by more proteins, especially effectors. The impact of other effectors on regulation of *RB*-mediated plant defense responses will be a focus of future research.

The prevalence of alternative splicing (AS) in plants not only contributes significantly to transcriptome, but also have function on proteome diversity. It is estimated that between 42% and 61% of all gene transcripts are alternatively spliced (Filichkin et al., 2010; Lu et al., 2010b; Marquez et al., 2012; Reddy et al., 2013; Klepikova et al., 2016). Several NLR genes undergo AS in the coding sequence or untranslated regions (UTRs; Yang et al., 2014; Lai and Eulgem, 2018). Our data indicate that the *RB* gene produces two different transcripts that encode either a full-length NLR protein that can confer resistance to *P. infestans*, or a truncated protein only containing the N-terminally domain of NLR. Premature termination codon (PTC)-containing transcripts that arise as a result of AS are often subjected to a nonsense-mediated mRNA decay (NMD) process that is intended to prevent their translation (Brogna and Wen, 2009). An analysis of transcripts produced during bacterial infection in Arabidopsis showed that the majority of CC (coiled coil)-NLR (65.1%) and TIR (Toll-like, Interleukin 1)-NLR (81.2%) transcripts exhibit evidence of NMD (Jung et al., 2020). However, transcripts containing PTCs can be used to express the truncated protein or have functional relevance (Liu et al., 2017; Nishimura et al., 2017). Our results indicate that the *RB* transcripts are still detectable by RT-PCR and the truncated RB_IR protein is detectable in plants (Fig. 1C), suggesting that the short transcripts are not immediately degraded. We further demonstrated the truncated protein RB_IR have an impact on immunity activation through the interaction with full-length RB protein. Previous evidence indicates that the CC domain of RB can self-oligomerize and this oligomerization is important in activation of immunity (Chen and Halterman, 2017; Zhao and Song, 2021). In this paper, we preliminarily demonstrated that overexpression of truncated RB_IR can disrupt the self-association of RB. The impact of this interaction on formation of a resistosome structure is therefore likely, but requires further confirmation. Such an investigation would help elucidate whether an RB_IR and RB_CDS hybrid structure is formed to prevent autoactivation due to RB_CDS self-oligomerization, thus avoiding inappropriate immune responses during regular plant growth.

For plants, homeostasis between growth and defense is vital for survival and reproduction in nature (He et al., 2022). Research on the role of *NLR* expression in regulating growth versus defense trade-offs in plants has been reported previously (Li et al., 2012; Tsuchiya and Eulgem, 2013; Cui et al., 2020; Huang et al., 2021; He et al., 2022). We found that the intron within the sequence of *RB* is alternatively spliced, this is a strategy to reduce elicitation of immune responses in the absence of an invading pathogen by regulating expression of the RB_CDS protein in the plant. Our findings suggest a strategy to reduce elicitation of immune responses in the absence of an invading pathogen by regulating expression of the functional plant *R* gene.

## Materials and methods

### RT-PCR, RT-qPCR analysis of *RB* splicing ratio and gene expression

Total RNA was extracted using E.Z.N.A.® Plant RNA Kit (R6827-01, OMEGA) following the manufacturer’s instructions and treated with RNase-Free DNase (M6101, Promega) to remove genomic DNA contamination. Approximately 700 ng RNA was used for reverse transcription with HiScript III 1st Strand cDNA Synthesis Kit (R312-01, Vazyme). RT-PCR analysis was performed with cDNA prepared from different potato or *N. benthamiana* leaf samples with gene specific primers (Supplemental Data Set S2). An equal amount of cDNA template in each reaction was verified by amplifying gene *EF1*α. The intensities of PCR products were quantified by ImageJ software. RT-qPCR was performed with 2µL of 3-fold-diluted cDNA in a total reaction volume of 20µL in technical duplicates using ChamQ Universal SYBR qPCR Master Mix (Q711-02, Vazyme Biotech) with an ABI 7500 Fast Real-time PCR system according to the manufacturer’s instructions. PCR conditions were as follows: 95°C for the 30s, followed by 40 cycles of 95°C for 5 s, 60°C for 34 s, and then 95°C for 15 s, 60°C for 1 min, 95°C for 30 s, 60 [for 15s for the melt curve. Relative transcript amount differences were calculated using the 2^-ΔΔCT^ method in ABI 7500 System Sequence Detection Software. Target-specific amplification efficiencies are given in Supplemental Data Set S4. Three biological replicates were performed and the results showed similar trends.

### Gene silencing and complementation in *N. benthamiana*

Virus-induced gene silencing (VIGS) was performed in *N. benthamiana* as previously described (Stam et al., 1997). For VIGS target design, the specific DNA fragments with 300 base pairs (bp) from GFP, *NbCWC15* (Niben101Scf04847g02012.1/Ni-ben101Scf00693g01009.1) predicted by VIGS tool (https://vigs.solgenomics.net/) were cloned into Tobacco Rattle Virus (TRV) construct. The sequences of the targets are listed in Supplemental Data Set S1. The constructs were introduced into *A. tumefaciens* GV3101 by electroporation. Suspensions of *A. tumefaciens* strain GV3101 harboring TRV1 and TRV2 constructs were mixed in a 1:1 ratio in infiltration buffer (10 mM MES, 10 mM MgCl_2_ pH 5.6, and 150 uM acetosyringone) to a final OD_600_ of 0.4, and then left at room temperature for 2h. TRV2-EV was used as a negative control. Two-week-old *N. benthamiana* plants were infiltrated with *A. tumefaciens* for VIGS assays, and their upper leaves were used 2-3 weeks later. The silencing efficiency was confirmed by RT-qPCR. For the complementation assays, the shuffled synonymous synthetic *StCWC15* sequences were generated by Sangon Biotech and cloned into expression constructs pK7WGF2 with a C-terminal GFP tag. Synthetic proteins were detected by western blot.

The 300-bp fragment of *NbPPS7a/b* (*Niben101Scf07798g01001.1*/*Niben101Scf01573g01003.1*), *NbSR30* (*Niben101Scf05298g04002.1*), *NbRS2Z32* (*Niben101Scf07227g00011.1*), *NbSGT1* (*Niben101Scf03241g00015.1*), *NbU1-70ka/b* (*NbS00011208g0006.1*/ *NbS00015731g0004.1*), *NbSF3b1a/b* (Niben101Scf06716g01016.1/ Niben101Scf06128g00004.1), *NbGRP7* (Niben101Scf00215g03004.1) were cloned and ligated into pK7GWIWG2D(II) to generate a hairpin silencing construct. The sequences of the targets are listed in Supplemental Data Set S1. The pK7GWIWG2D(II) empty vector was used as a control. Six-to seven-week *N. benthamiana* leaves were infiltrated with GV3101 *A. tumefaciens* strains containing either the silencing construct or empty vector control diluted to an OD_600_=0.5.

### Growth of plant material and microbial strains

*Solanum tuberosum* cv. Favorita and transgenic plants were grown on Murashige and Skoog (MS) medium in a growth chamber under 16 h light/8 h dark conditions at a constant temperature of 24 °C and light intensity of 100 μM m^−2^ s^−1^. Plants in soil were growth with 12 h light/12 h dark conditions at a constant temperature of 22°C and light intensity of 153 μM m^−2^ s^−1^.

*N. benthamiana* plants were grown and maintained in the greenhouse with 16 h light at 25°C and 8 h dark at 20°C throughout the experiments. *P. infestans* strains JH19 (Howard et al., 2013), and 88069^td^, expressing the red fluorescent marker tandem dimer RFP (tdTomato) (Dagdas et al., 2016) were grown on rye sucrose agar (RSA) at 18°C. *P. capsici* strain LT1534 was grown on V8 medium at 25[. *Escherichia coli* strains DH5α, JM109, and BL21 and *A. tumefaciens* strain GV3101 were grown on Luria-Bertani (LB) medium at 37 °C and 30 °C, respectively.

*S. bulbocastanum* accession PT29 (PI 243510) was originally obtained from the United States Potato Genebank (NRSP-6) in Sturgeon Bay, WI. The accession sto389-4 containing native promoter *Rpi-sto1*, blb8005 containing native promoter *RB* were kindly provided by Prof. Jack H. Vossen. Binary plasmids pBINPLUS: *Rpi-sto1*, pBINPLUS: *RB* were constructed in house and transformed into cv Desiree using Agrobacterium mediated transformation (Zhu et al., 2015).

### Plasmid construction

All primers and plasmids used for cloning and experiments are listed in Supplemental Data Set S2. All the templates used for PCR amplification come from genomic DNA (gDNA) or complementary DNA (cDNA) of *P. infestans* (strain T30-4), *N. benthamiana*, and *Solanum tuberosum* cv. Favorita using 2 × Phanta Max Master Mix (P515-01, Vazyme Biotech). FastPure Plasmid Mini Kit, FastPure Gel DNA Extraction Mini Kit and ClonExpress Ultra One Step Cloning Kit (Cat No. DC201-01, DC301-01 and C115-01) were used for nucleic acid purification, digestion, and ligation.

### Co-immunoprecipitation assays and GST pull-down

The *IPI-O1* gene without the signal peptide was inserted into pK7WGF2 with N-terminal RFP tag using ClonExpress Ultra One Step Cloning Kit (C115-01, Vazyme Biotech), and *StCWC15a/b* were inserted into pK7WGF2 with C-terminal GFP for expression in *N. benthamiana.* Leaves of 6-week-old *N. benthamiana* plants were co-infiltrated with RFP-tagged *IPI-O1*, GFP-tagged *StCWC15a/b*, and the P19 silencing suppressor. Two days after agro-infiltration, the leaves were frozen in liquid nitrogen and ground to powder using a mortar and pestle. Proteins were extracted with extraction buffer (10% [vol/vol] glycerol, 25mM Tris pH=7.5, 1mM EDTA, 150mM NaCl, 2% [wt/vol] PVPP, 10mM DTT, 1×Protease Inhibitor Cocktail (Sigma-Aldrich), 0.1% Tween 20). The samples were centrifuged at 4°C for 20 min at 12,000 × g, and the supernatant was transferred to a new tube. For GFP-IP, 1 mL of supernatant was incubated at 4°C for 2 h with GFP-Trap_A beads (Chromotek, Planegg-Martinsried, Germany). The beads were collected by centrifugation at 800 × g and washed four times in 1 mL washing buffer (25 mM Tris pH=7.5, 150 mM NaCl and 1mM EDTA, 10% [vol/vol] glycerol, 0.1% Tween 20). The beads were boiled for 10 min at 95°C. SDS-PAGE was performed with the supernatant, and co-IP of StCWC15a/b with IPI-O1 was detected using anti-GFP or anti-RFP (Chromotek, Planegg-Martinsried, Germany).

For *in vitro* GST pull-down assays, GST-StCWC15a/b and His-IPI-O1 were expressed in *E. coli* strain BL21. Co-precipitation of StCWC15a/b with IPI-O1 was examined by protein blotting before (input) and after affinity purification (pull-down) using glutathione agarose beads (GE Healthcare). Interactions were detected using anti-GST and anti-His (Abmart).

### P. infestans and P. capsici infection assays

*P. infestans* strains were grown on RSA medium plate at 18°C for 14 d. The sporangia were scraped into cold, distilled water and incubated at 4°C for 1-2 h until zoospore release. The zoospore suspension was adjusted to 100 zoospores/µL before inoculation. Ten microliter drops of *P. infestans* zoospores were placed on the abaxial side of detached leaves. The inoculated leaves were kept at 18°C for 4-7 days. The whole potato infection assays used a zoospore suspension sprayed on the plant followed by incubation in a controlled environment chamber at 20°C for 6 days. The infected *N. benthamiana* leaves were stained with trypan blue. Leaves were soaked in trypan blue solution (10 mL 85% lactic acid aqueous, 10 mL water-saturated phenol, 10 mL 98% glycerol, 10 mL distilled water, 15 mg trypan blue) overnight. Stained leaves were then transferred into ethanol and incubated at 37°C for 24 h before observation. *P. capsici* strains were grown on V8 medium plate at 25 [for 3 d. Ten pieces of mycelium with a size of about 2 mm^2^ were cultured in V8 liquid medium for 4 days. The culture medium was discarded, the mycelium was washed with distilled water three times, and distilled water was added with just enough to cover the mycelium. The culture was incubated at 25 [for 12 hours until sporangia were observed, then was placed at 4°C for 20 minutes to stimulate zoospores release.

### Agroinfiltrations

Construct plasmids were transformed into *A. tumefaciens* strain GV3101 and grown overnight in Luria-Bertani (LB) culture medium with the antibiotic kanamycin and rifampicin at 30°C. Bacteria were harvested by centrifugation at 5000g for 4 min and then resuspended and diluted in an agroinfiltration buffer (10 mM MgCl_2_, 150 mM acetosyringone, 10 mM MES) to an optical density of OD_600_ = 0.01-1.0 depending on the experiments. After 2-4 h of incubation at room temperature, they were infiltrated with a needleless syringe into the abaxial side of leaves from 4–5-week-old *N. benthamiana* plants. For the co-expression of multiple constructs, suspensions carrying each construct were thoroughly mixed before infiltration.

### Confocal microscopy and bimolecular fluorescence complementation (BiFC) assays

Confocal microscopy was performed on *N. benthamiana* leaf patches post infiltration by LSM 980 laser-scanning microscope with 20×/0.8 or 60×/0.95 objective lens (ZEISS LSM 980 with Airyscan2). Parameters that are used for fluorescence detection are GFP/YFP, (excitation 488 nm, emission 505–540 nm), RFP (excitation 514 nm, emission 518–582 nm), DAPI (excitation 405 nm, emission 430–480 nm). Leaves were stained with 2 μg/mL of 4’, 6-diamidino-2-phenylindole (DAPI) for 5 min to indicate nuclei. ZEISS ZEN Microscope Software was used to process the images.

pGTQL1221YC and pGTQL1211YN (nYFP/cYFP) vectors were used for bimolecular fluorescence complementation (BiFC) assays (Lu et al., 2010a). Coding regions of *IPI-O1* and *IPI-O1^nls1^* without a signal peptide and *StCWC15a/b* were amplified with primers containing attB tails and BP-cloned into pDONR207 (Invitrogen). To generate BiFC constructs, constructs were integrated into destination vectors pGTQL1221YC or pGTQL1211YN by LR recombination using Gateway LR clonase II (Invitrogen). The constructs were transformed into *A. tumefaciens* strain GV3101. Two days after agroinfiltration, the Zeiss 980 laser scanning confocal microscope was used for YFP fluorescence detection.

### Transformation and RNAi of potato and *N. benthamiana*

For generating *RB* transgenic potato and transgenic *N. benthamiana*, the entire *RB* gene sequence (3592 bp), *RB_CDS* transcript (2913 bp), *RB_IR* transcript (507 bp) and *RB_IVS* mutant were cloned into pBinGFP2 binary vector containing the *CaMV35S* promoter. The constructs were transformed into the *A. tumefaciens* strain GV3101. The copy number of transgenic potato lines is calculated as reported (Yang et al., 2005). To generate stable transgenic potato plants, stems were used for transformation. Stem segments of *Solanum tuberosum* cv. Favorita were incubated in MS-medium (Murashige and Skoog, 1962). Agrobacterium containing each plasmid was grown and diluted to a cell concentration of OD_600_=0.5. Stem segments were added and incubated in the dark for 45 mins. After co-incubation, stem segments were transferred to MS solid medium supplemented with 2.5 mg/L zeantinribose, 0.2 mg/L NAA, 0.02 mg/L GA_3_ for 48 hours. After that, stems were washed with 300mg/L Timentin and transferred to MS solid medium supplemented with 2.5 mg/L zeantinribose, 0.2 mg/L NAA, 0.02 mg/L GA_3_ and 200 mg/L Timentin for 2 weeks. For callus induction, stems segments were kept on MS solid medium with 2 mg/L zeantinribose, 0.02 mg/L NAA, 0.02 mg/L GA_3_, 300 mg/L Timentin and 50 mg/L kanamycin. Every 14 days, stem segments or developing calli were placed onto new medium. After bud initiation formation (following 4-5 transfers), shoots were cut off from the callus and transferred to MS-medium containing 300 mg/L Timentin and 50 mg/L kanamycin. The *N. benthamiana* stable transformation was conducted according to a previously established protocol (Davarpanah et al., 2009).

For hairpin RNA interference (RNAi), the chalcone synthase (CHSA) intron was amplified from plasmid pFGC5941. The PCR-amplified 3′StCWC15 cDNA fragments overlapped the 5′ and 3′ end of the CHSA intron from two directions. The overlapped fragment was inserted in the p1300 vector and transformed into *A. tumefaciens* strain GV3101 before potato transformation. Transformation is the same as potato stable overexpression transformation. *StCWC15s* silencing transgenic lines were confirmed by RT-qPCR in buds using the primers indicated in Supplemental Data Set S2.

### 5’ RACE

Rapid amplification of cDNA ends (5’ RACE) was used to determine the transcripts of the *RB* gene using the SMARTer RACE cDNA Amplification Kit (Clontech) according to the protocol. For synthesis of first-strand cDNA, we designed the *RB* specific primer (Supplemental Data Set S2). Next, we used 10 μM *RB* specific primer and the provided UPM primer in the kit for RACE PCR. Amplification was performed using a touchdown PCR protocol with five cycles of 94°C for 30 s, 72°C for 4 min, then five cycles of 94°C for 30 s, 70°C for 30s, 72°C for 4 min, final 25 cycles of 94°C for 30 s, 68°C for 30 s, 72°C for 4 min. The obtained 5′RACE products were gel excised, introduced into the pUC19 vector using in-fusion cloning, and sequenced by Sanger sequencing to detect the transcripts.

### LUC/GUS reporter assays

For LUC/GUS reporter system assays in *N. benthamiana* leaves, a 1986-bp *RB* fragment including the intron structure was amplified from *RB* genomic DNA and fused with the *LUC* gene, followed by ligattion into pCAMBIA1300 vector behind the CaMV 35S promoter. The *GUS* gene was ligated into pCAMBIA1300 vector behind the CaMV 35S promoter. Four- or five-week-old *N. benthamiana* leaves were co-infiltrated with Agrobacterium strain GV3101 harboring the RB-LUC and GUS constructs resuspended in infiltration buffer. At 48 h after infiltration, two samples from each infiltrated area were collected. One of the samples was sprayed with 1 mM D-luciferin substrate and incubated for 5 min in the dark. For quantification of LUC activity, *N. benthamiana* leaf discs were collected at 48 h after infiltration (hpi) and incubated with 1 mM D-luciferin in a 96-well plate, and LUC activity was quantified with the Promega GloMax Navigator microplate luminometer (Promega, USA). The other sample was used for extracting GUS protein. For quantification of GUS activity, extracted protein was incubated with 1mM MUG (4-methylumbelliferyl β-D-glucuronide) (Sigma)substrate for 30 min at 37[. The GUS fluorescence intensity was quantified with the Promega GloMax Navigator microplate luminometer (Promega, USA). The pCAMBIA1300-GUS was used as the internal control. The LUC images were photographed using a Tanon-5200 Multi Chemiluminescent Imaging System (Tanon, China).

### Yeast two-hybrid screening

To identify interacting partners with the *P. infestans* effector IPI-O1, a yeast two-hybrid interaction screen was performed by Hybrigenics Services (Évry-Courouronnes, France) using IPI-O1 as a bait in an ULTImate Y2H library constructed from *N. benthamiana*. IPI-O1, encoding amino acids 2-132, was cloned into the inducible pB35 vector as a fusion with GAL4 at the amino terminus. The prey library was prepared in the proprietary pP6 Hybrigenics vector. A total of 81.6 million interactions were analyzed and 254 clones were processed by sequencing of the prey inserts. These prey fragments were fused out-of-frame with the Gal4 activation domain (AD) in pP6. However, *Saccharaomyces cerevisiae* are able to translate a percentage of the correct fusion proteins using ribosomal frameshift (Fromont-Racine et al., 1997; Vidal and Legrain, 1999). For this reason, it is possible to identify true interacting proteins from out-of-frame clones.

### Hypersensitive response phenotype and ion leakage

Hypersensitive cell death during the ETI process was scored at 4 dpi and was quantified using an ion leakage assay (Mittler et al., 1999). Membrane ion leakage was determined by measuring electrolytes leaked from leaf discs. For each measurement, five 9 mm leaf discs were immersed in 5 mL distilled water at room temperature (22-25°C) for 3h. After which, the initial conductivity was measured with Multi-parameter tester (Mettler Toledo S400-K), referred to as value A. Total conductivity was determined after boiling at 95°C for 25 min and cooling to room temperature; this is referred to as value B. The ion leakage was expressed as the percentage of the initial conductivity versus the total conductivity (value A/value B) ×100%.

### Salicylic acid quantification

For SA quantification, three replicates of each genotype transgenic potato were collected. Approximately 100 mg (fresh weight) of leaf tissues were frozen in liquid nitrogen, ground and extracted with 1 mL 80% methanol (Anaqua™ Chemicals Supply, CAS: 67-56-1) containing 1% formic acid (Anaqua™ Chemicals Supply, CAS: 64-18-6) mixture. The resulting solution was then vortexed and placed in a shaker at 4□ for 24 h. Samples were centrifuged 10 min at 15,000 rpm, and supernatant was transferred to new tubes. Add 1 mL mixture and 0.3 g PVPP in the precipitate to shaker at 4 [for 24 h again. Mix two times supernatant and drying most of the liquid with a vacuum pump. Using 200 μL of 80% methanol for redissolve. Supernatants were filtered through 0.2 mm PTFE membrane (Millipore, Bedford, MA) and transferred to auto sampler vials. Quantification was performed by HPLC with the ACQUITY UPLC HSS C18 column.

### Inhibitor assay

Pladienolide B (CAS:445493-23-2) was purchased from Adooq. PB, dissolved in DMSO, was infiltrated into *RB* transgenic *N. benthamiana*. The exact concentration of DMSO was used as control infiltrated in *N. benthamiana*.

Lanthanum (III) chloride (LaCl_3_) (Sigma, CAS: 10099-58-8) was dissolved in water and infiltrated at a concentration of 2 mM.

### Protein sequence alignments

Peptide sequences of CWC15 homologs were extracted from published potato pan-genome repositories (Tang et al., 2022). All selected protein sequences were aligned by R-msa packages (Version 1.33.0) using ClustalW, and visualized

## ACKNOWLEDGMENTS.

We thank Sebastian Schornack (University of Cambridge), Jiamu Du, Zhe Wu (Southern University of Science and Technology), Jiming Jiang (Michigan State University), Yan Wang (Nanjing Agriculture University) for helpful discussions. The author responsible for distribution of materials integral to the findings presented in this article in accordance with the policy described in the Instructions for Authors (https://academic.oup.com/plcell/pages/General-Instructions) is: Suomeng Dong. This work was supported by the Chinese National Science Funds Grants (32130088, 32102234) and the China Agriculture Research System-potato (CARS-potato, CARS-09-P20). This research was supported in part by the United States Department of Agriculture (USDA) Agricultural Research Service (ARS). Funding was provided through USDA National Institute of Food and Agriculture (NIFA) Plant-Associated Microbes and Plant-Microbe Interactions award number 2014-67013-21593. The findings and conclusions in this publication are those of the author(s) and should not be construed to represent any official USDA or United States Government determination or policy.

## Notes

### Competing Interest Statement

The authors have declared no competing interest.

### Summary of Updates

Revisions to address reviewer comments. Correction to fix an incorrect image used in Figure2.

