## Supplemental Fig. for "Alternative splicing of a potato disease resistance gene maintains homeostasis between development and immunity"

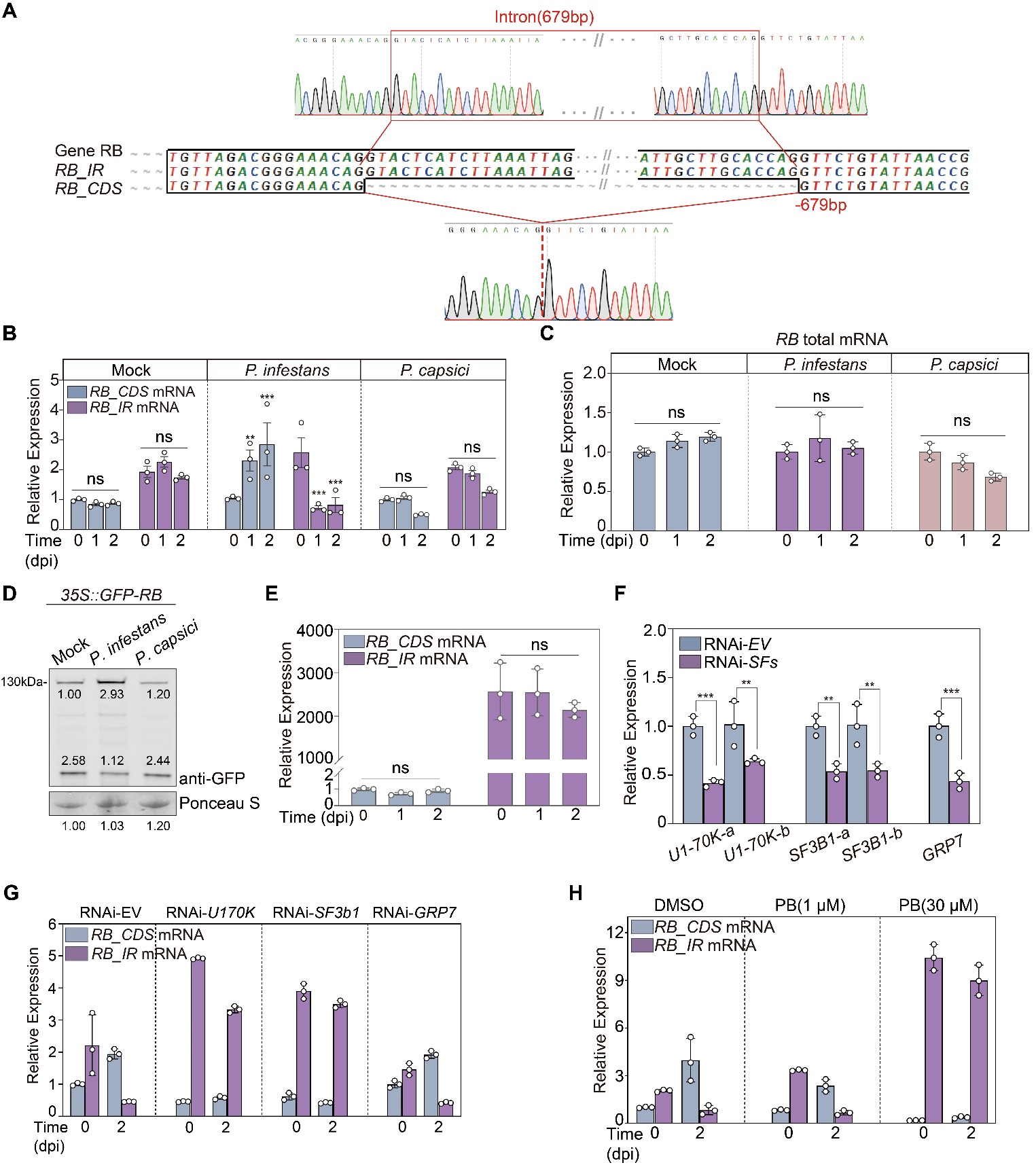

**Supplemental Figure S1**. ***RB* undergoes AS and produces two transcript isoforms.**

**(A**) Sanger sequencing traces of intron junction regions confirming RT-PCR products in Fig. 1B. **(B**) Transcript levels of *RB_CDS* and *RB_IR* mRNAs in *RB* transgenic potato following inoculation with *P. infestans*. Quantitative PCR with reverse transcription (RT–qPCR) analysis was performed using isoform-specific primers (Supplemental Data Set S2) to measure the transcript levels of two isoforms. Bars represent the mean ± SD from three independent replicates. **(C)** Total *RB* transcription level was performed using specific primer (Supplemental Data Set S2) located in *RB* exon1 by RT–qPCR. Bars represent the mean ± SD from three independent replicates. **(D)** RB_CDS and RB_IR protein expression following *P. infestans* and *P. capsici* inoculation. *N. benthamiana* leaves were infiltrated with Agrobacterium carrying *35S::GFP-RB*. The infiltrated areas of leaves were inoculated with water (mock), *P. infestans*, *P. capsici* zoospores at one day post inoculation. Leaf tissues were sampled and used for protein extraction one day post pathogen inoculation (dpi). The GFP-RB protein was enriched using anti-GFP beads before detection using anti-GFP antibody. Protein loading was visualized by Ponceau S staining. The intensities under the signal bands of each protein were determined using ImageJ software. The above experiments were repeated 3 times with similar results. **(E)** Transcript levels of *RB_CDS* and *RB_IR* mRNAs in splicing mutant *RB_IVS* following inoculation with *P. infestans*. RT–qPCR was performed using isoform-specific primers to measure the transcript levels of two isoforms. Bars represent the mean ± SD from three independent replicates. **(F)** RT–qPCR demonstrating the silencing efficiency of RNAi constructs targeting splicing factors (SFs). Transcript levels of each gene were calculated relative to the RNAi-*EV* control. Bars represent the mean ± SD from three independent replicates. For all graphs, ns = no significant difference, two asterisks denote a statistical difference at P<0.01, three asterisks denote a statistical difference at P<0.001 using a two-way ANOVA with Tukey’s test. **(G)** Detection of *RB_CDS* and *RB_IR* mRNAs following agroinfiltration of the indicated RNAi-hairpin constructs, and inoculation with *P. infestans* zoospores one day after infiltration. RNA was extracted 2 days after pathogen inoculation. RT-qPCR analysis was performed using isoform-specific primers to measure the transcript levels of the two isoforms. Bars represent the mean ± SD from three independent replicates. **(H)** Transcript levels of *RB_CDS* and *RB_IR* mRNAs in *RB* transgenic *N. benthamiana* following treatment with DMSO (control), 1 µM Pla-B, or 30 µM Pla-B and inoculation with *P. infestans* zoospores at 2 dpi by RT-qPCR. Bars represent the mean ± SD from three independent replicates.

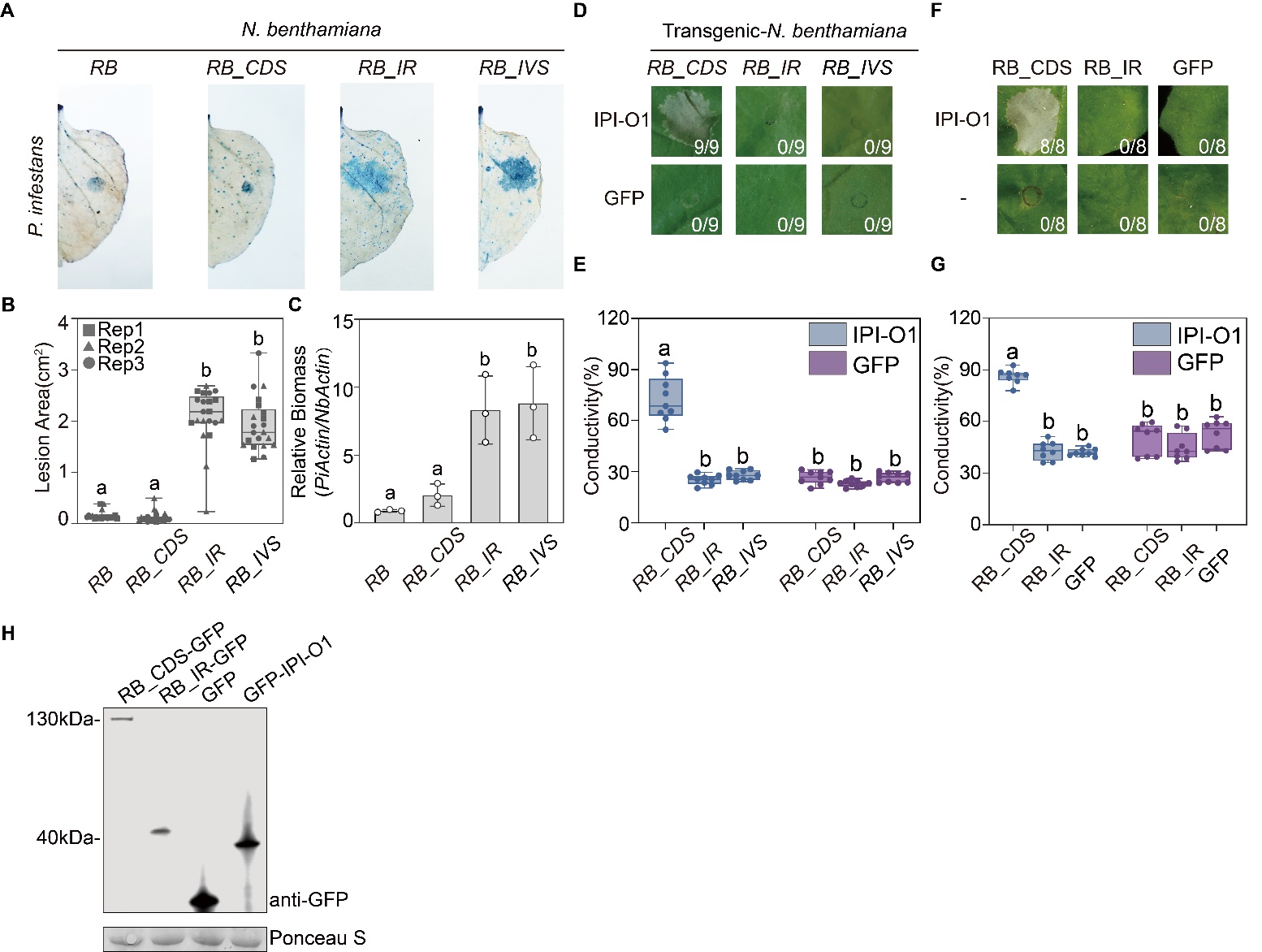

**Supplemental Figure S2**. **Intron retention in *RB* results in susceptibility to *P. infestans.***
**(A)** Zoospores of *P. infestans* strain JH19 were inoculated on detached transgenic *N. benthamiana* leaves expressing the indicated gene variants. Leaves were stained with trypan blue and photographed at 5 dpi. **(B)** Lesion areas(cm^2^) of inoculated leaves in (A) were calculated using ImageJ. Data represent the mean ± SD from 21 replicates from three independent experimental replicates. Different letters above boxes denote a significant difference at P<0.001 using one-way ANOVA with Tukey’s HSD test. Different shapes indicate biological replicates. **(C)** Relative biomass of *P. infestans* was determined by qPCR of *P. infestans* genomic DNA normalized to potato genomic DNA at 5 dpi. Bars represent the mean ± SD from three independent replicates. Different letters above bars denote a significant difference at P<0.01 using one-way ANOVA with Tukey’s HSD test. **(D)** Indicated transgenic *N. benthamiana* leaves were agroinfiltrated with IPI-O1 or GFP. Cell death was visually assessed and photographed at 4 dpi. The numbers represent the number of hypersensitive cell death responses compared to all infiltration points. **(E)** Results of electrolyte leakage assays of infiltrated areas. Data represent the mean ± SD from 9 replicates. Different letters above each bar represent a significant difference at P<0.001 using two-way ANOVA with Tukey’s HSD test. **(F)** Indicated *RB* constructs and GFP were agroinfiltrated alone or with IPI-O1. Cell death was visually assessed and photographed at 4 dpi. The numbers represent the number of hypersensitive cell death responses compared to all infiltration points. **(G)** Results of electrolyte leakage assays of infiltrated areas. Data represent the mean ± SD from 8 replicates. Different letters above each bar represent a significant difference at P<0.001 using two-way ANOVA with Tukey’s HSD test. **(H)** Expression of GFP-tagged fusions of RB_CDS, RB_IR and IPI-O1 in *N. benthamiana* from part (F) was confirmed by protein blot using a GFP antibody. The GFP-RB_CDS/RB_IR were enriched using anti-GFP beads before detection. Protein loading is visualized by Ponceau S staining.

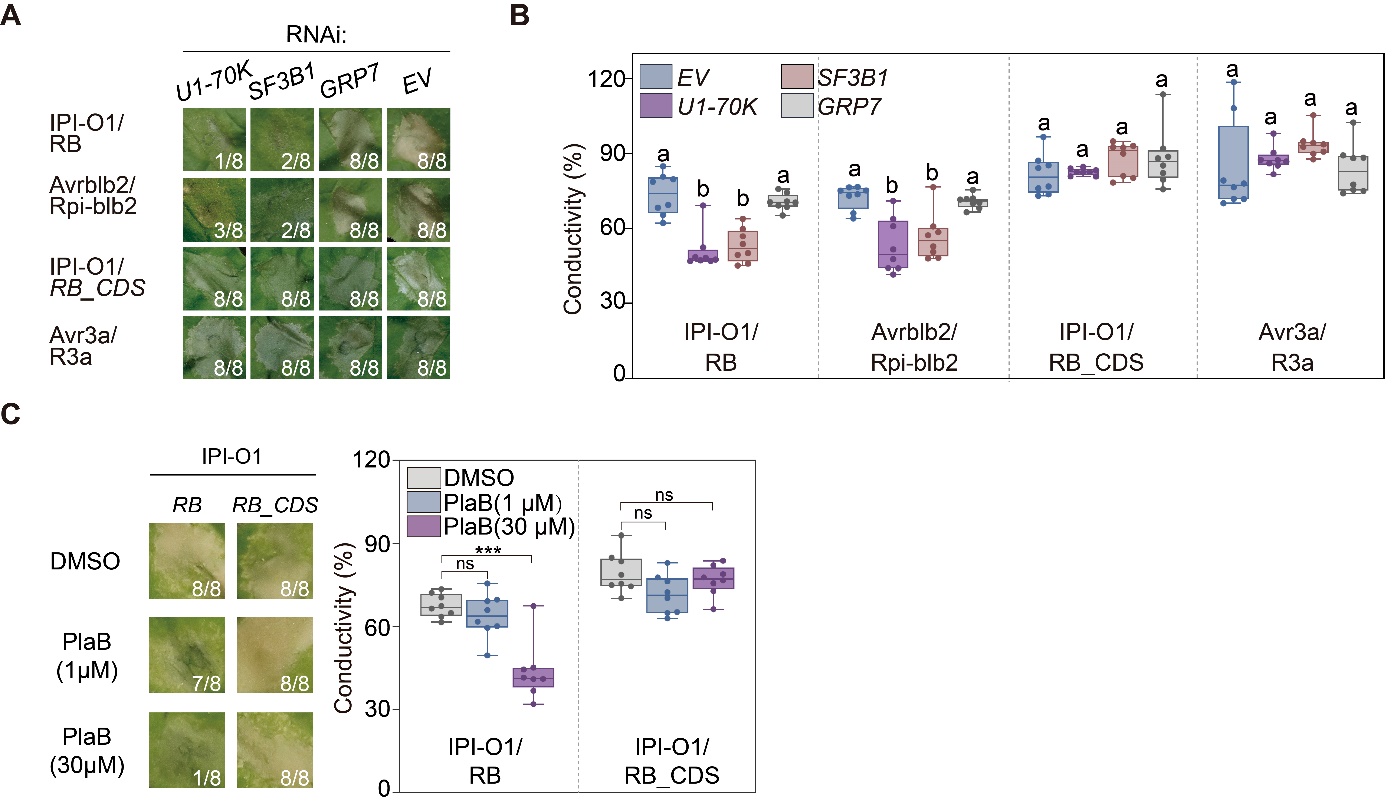

**Supplemental Figure S3**. **Spliceosome activity is necessary for *RB* induced HR.**
**(A)** Agroinfiltration of the indicated RNAi-hairpin constructs to silence *U1-70K*, *SF3B1* and *GRP7* in *N. benthamiana*, followed by co-agroinfiltration of the indicated Avr/R gene combinations. Cell death was visually assessed and photographed at 4 dpi. The numbers represent incidence ratios of hypersensitive cell death responses. **(B)** Electrolyte leakage assay of cell death. Data represent the mean ± SD from 8 replicates. Different letters above each bar represent a significant difference at P<0.01 using two-way ANOVA with Tukey’s HSD test. **(C)** Agroinfiltration of IPI-O1 in *RB* and *RB_CDS* transgenic *N. benthamiana* with DMSO (control), 1 μM splicing modulator Pladienolide B (PB), or 30 μM PB. Cell death was visually assessed and photographed at 4 dpi (left). Numbers show the ratio of infiltrated areas with cell death compared to all infiltrated areas. Electrolyte leakage was monitored by conductivity measurements following the same treatments (right). Data represent the mean ± SD from 8 replicates. ns = no significant difference, three asterisks denote a statistical difference at P<0.01 using a two-way ANOVA with Tukey’s test.

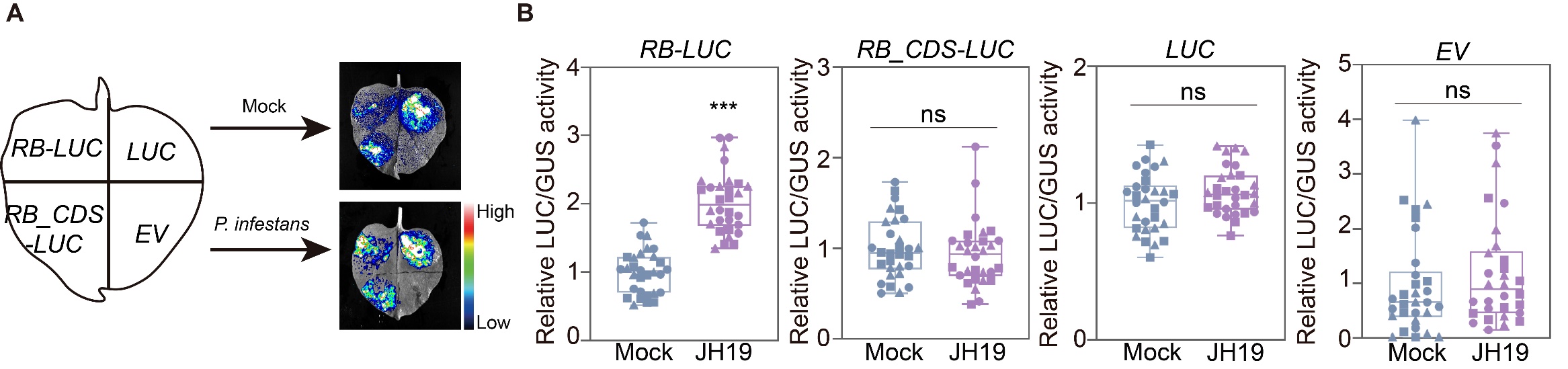

**Supplemental Figure S4. Luciferase assay to monitor *RB* intron splicing following *P. infestans* inoculation.**
**(A)** Representative photo of a *N. benthamiana* leaf infiltrated with agrobacterium carrying (clockwise) *RB*-*LUC*, *LUC*, empty vector (EV) control, or *RB_CDS-LUC*. One day after agroinfiltration, the infiltrated regions were treated with *P. infestans* or water (mock). The photo was taken at 1 dpi. **(B)** The activity of *RB*-*LUC, RB_CDS-LUC, LUC* and EV was normalized to GUS activity. The results represent 30 individual samples isolated from leaves agroinfiltrated with each indicated construct and GUS, followed by treatment with *P. infestans* or water. The LUC/GUS activities were relative to the activity of mock treatment. The data represent the mean ± SD from 30 individual samples from 3 experimental replicates. The different shapes of the data indicate independent biological replicates. ns=no significant difference and three asterisks denote a statistical difference at P<0.01 using two-sided Welch’s t-test.

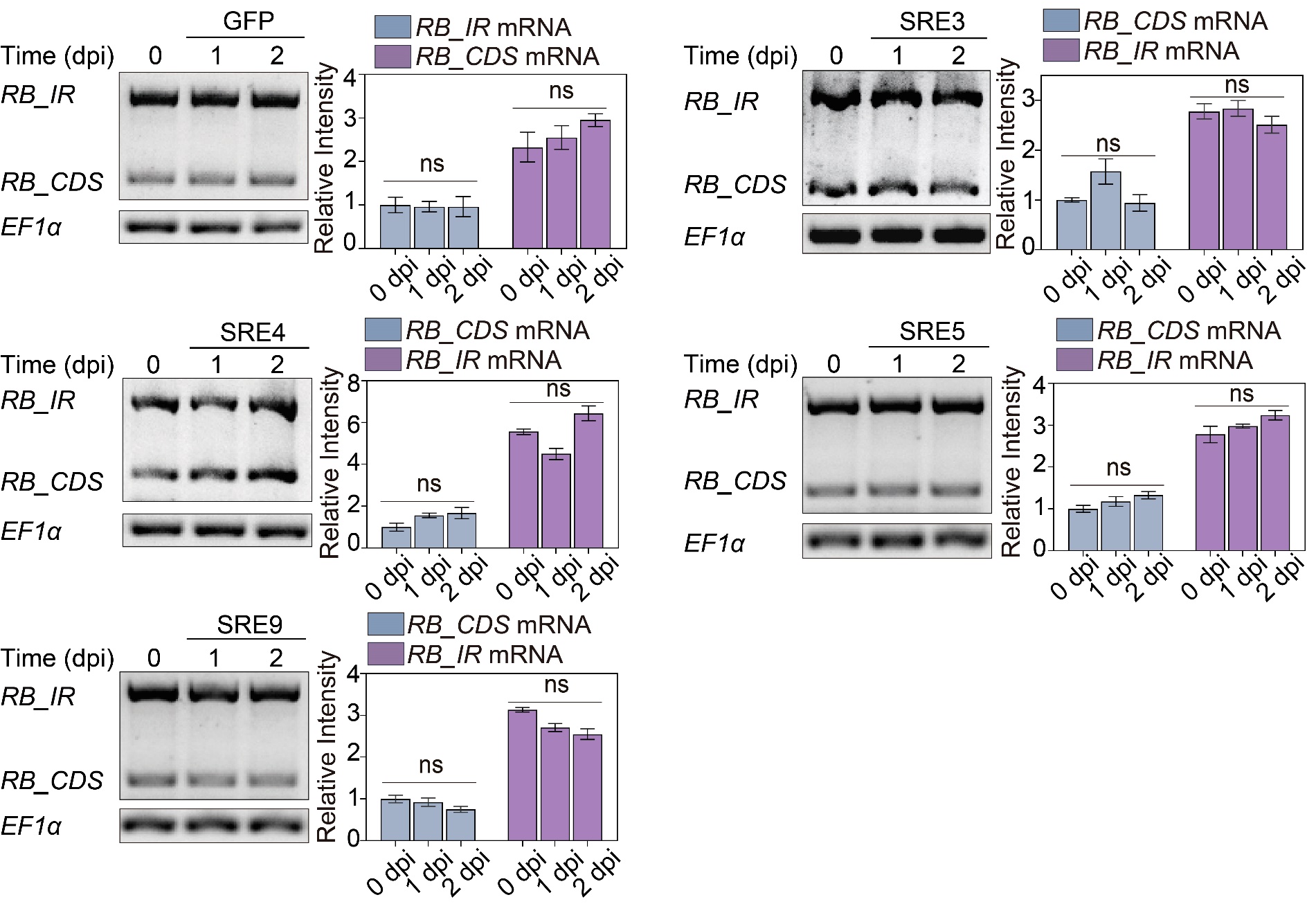

**Supplemental Figure S5. IPI-O1 and its homologue can modulate *RB* AS change and activate *RB*-mediated hypersensitive response**

On the left, transcript levels of *RB_IR* and *RB_CDS* were determined by RT-PCR following agroinfiltration of SRE proteins into leaves of *RB* transgenic *N. benthamiana*. On the right, data represent relative intensity of RT-PCR products normalized with internal control *NbEF1α*. Intensities of the bands were quantified using ImageJ software, and the relative abundances of *RB* isoforms are relative to *RB_CDS* at 0 dpi. Bars represent the mean ± SD from three independent replicates. ns = no significant difference using two-way ANOVA with Tukey’s test.

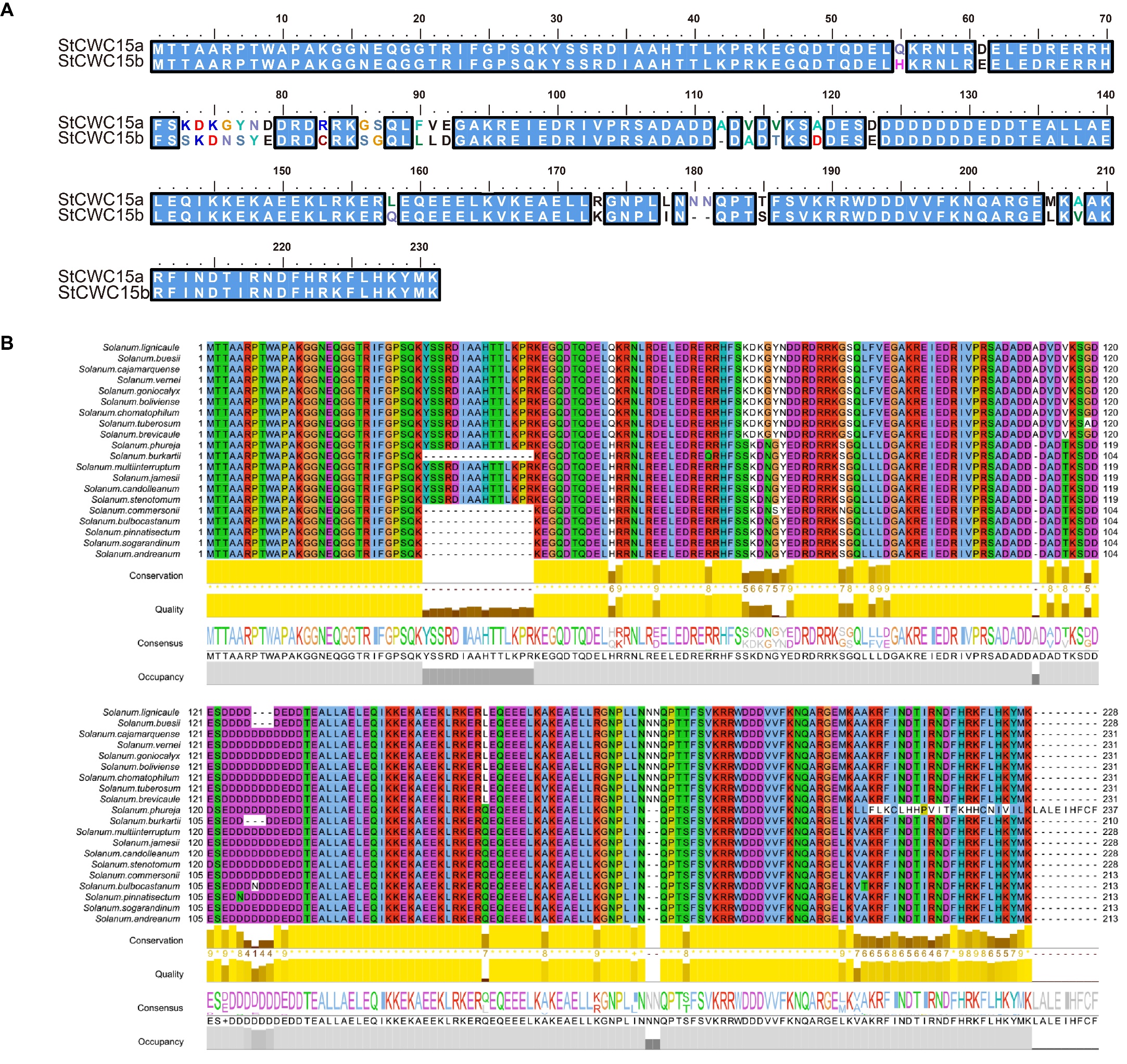

**Supplemental Figure S6. The splicing factor CWC15 interacts with IPI-O1 and is conserved in potato.**
**(A)** Alignment comparing CWC15a and CWC15b in double-monoploid potato DM 1-3. **(B)** Alignment comparing several CWC15 homologs in different potato cultivars and wild species. Individual amino acids are depicted in color scheme, degree of overall amino acid conservation is shown both as line plot and sequence logo.

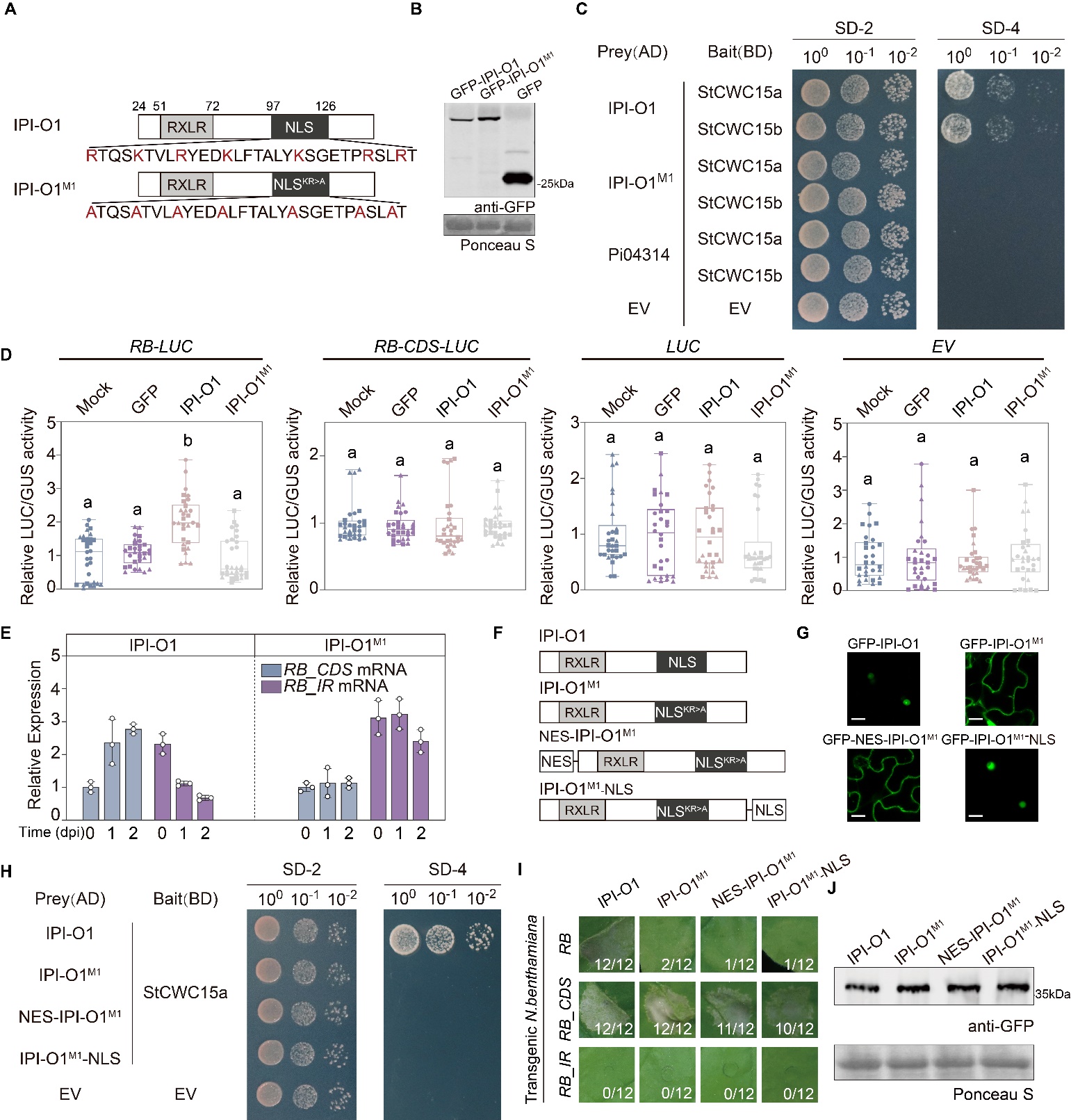

**Supplemental Figure S7. The splicing mutant IPI-O1^M1^ cannot interact with CWC15 and modulate *RB* AS.**

**(A)** Schematic diagrams show the alanine replacements within IPI-O1^M1^. **(B)** Expression of GFP-tagged fusions of IPI-O1 and IPI-O1^M1^ in *N. benthamiana* was confirmed by protein blotting using an anti-GFP antibody. Equal protein loading was visualized by Ponceau S staining. **(C)** IPI-O1, but not IPI-O1^M1^, can interact with CWC15 homologs in potato. Yeast transformants were spotted as 10-fold serial dilutions on SD medium without leucine and tryptophan (SD–2), or without histidine, leucine, adenine, and tryptophan (SD–4). Yeast cells were photographed 2-3 days later. BD and AD are bait and prey EVs, respectively. Experiments were performed at least three times with similar results. **(D)** Results of co-expression of *RB-LUC* reporters with mock, GFP, IPI-O1 and IPI-O1^M1^ in *N. benthamiana* leaves by agroinfiltration. The LUC/GUS activity of *RB*-*LUC, RB_CDS-LUC, LUC* and EV of infiltrated samples was calculated at 2 dpi. The LUC/GUS activities were relative to the activity of each construct with mock treatment. Data represent the mean ± SD of 30 replicates. Different letters above each bar represent a significant difference at P<0.001 using one-way ANOVA with Tukey’s HSD test. **(E)** The relative expression level of *RB* transcripts following overexpression of IPI-O1 and IPI-O1^M1^ at 0, 1, and 2 days post infiltration (dpi) by RT-qPCR. Bars represent the mean ± SD from three independent replicates. **(F)** Schematic diagrams of IPI-O1, IPI-O1^M1^, and IPI-O1^M1^ fused to NLS or NES. **(G)** GFP-IPI-O1, GFP-IPI-O1^M1^, GFP-NES-IPI-O1^M1^ and GFP-IPI-O1^M1^-NLS were expressed in *N. benthamiana* leaves for 48 h before imaging. Scale bar = 20 µm. **(H)** Yeast transformants were spotted as 10-fold serial dilutions on SD medium without leucine and tryptophan (SD–2), or without histidine, leucine, adenine, and tryptophan (SD–4). Yeast cells were photographed 2-3 days later. BD and AD are bait and prey EVs, respectively. Experiments were performed at least three times with similar results. **(I)** Visual examples and the number of hypersensitive cell death responses compared to all infiltration points following agroinfiltration of IPI-O1, IPI-O1^M1^, NES-IPI-O1^M1^, and IPI-O1^M1^-NLS in leaves of *RB*, *RB_CDS*, and *RB_IR* transgenic *N. benthamiana*. Cell death was visually assessed and photographed at 4 dpi. **(J)** Expression of GFP-tagged fusions in *N. benthamiana* was confirmed by protein blotting using an anti-GFP antibody. Protein loading was visualized by Ponceau S staining.

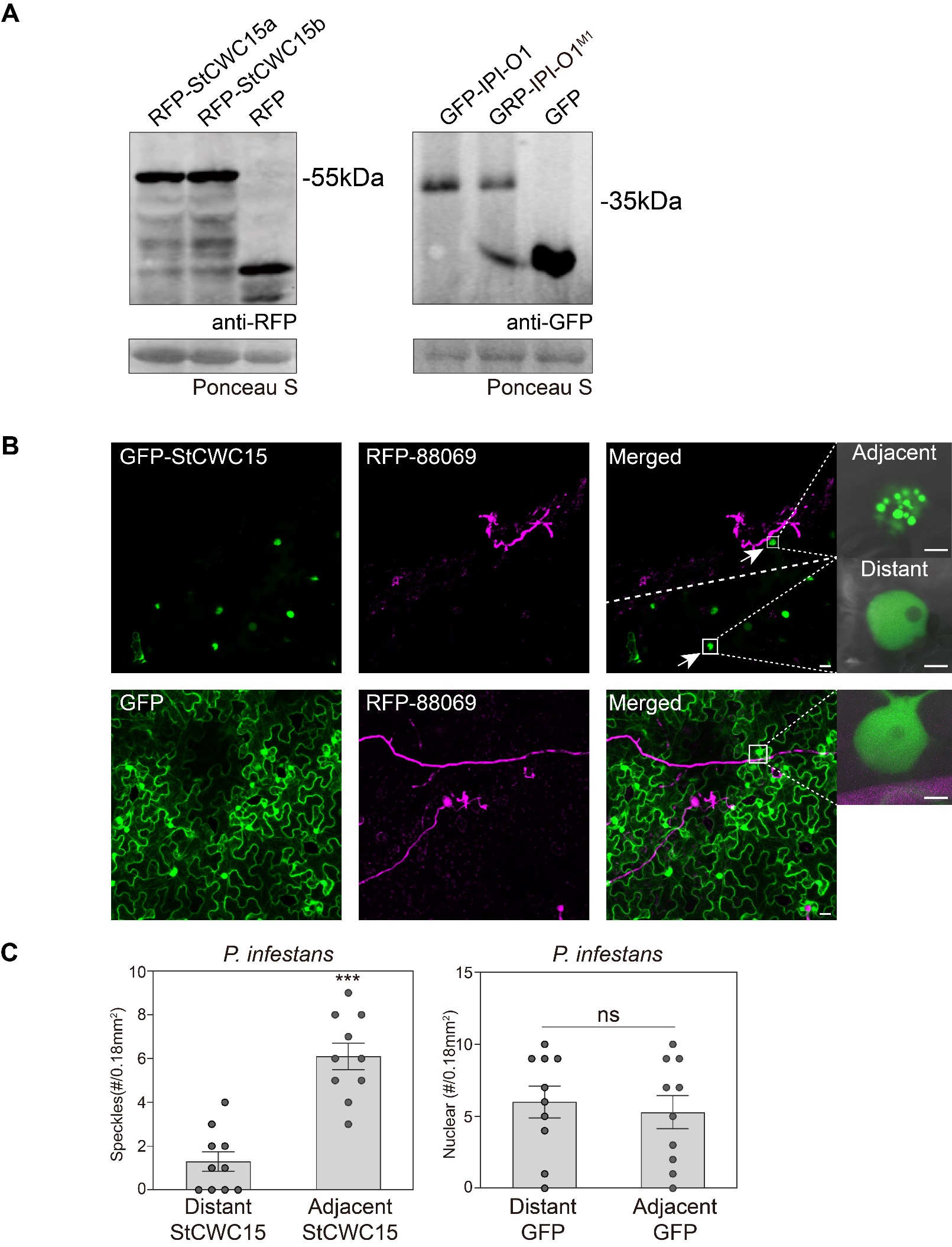

**Supplemental Figure S8. StCWC15 is re-localized to nuclear speckles during *P. infestans* infection.**

**(A)** Immunoblot analysis of protein in the co-localization experiment. RFP tagged fusions with StCWC15a/b were confirmed by protein blotting using an anti-RFP antibody. Protein loading is visualized by Ponceau S staining. GFP tagged fusions with IPI-O1, IPI-O1^M1^, and GFP proteins were confirmed by immunoblotting using an anti-GFP antibody. **(B)** Images are representative of the different patterns observed in GFP–StCWC15 re-localization from the nucleoplasm to speckles. Nucleolar GFP–StCWC15 fluorescence near the haustoriated cells was detected in speckles (upper panel) while in cells distant from the haustoriated cells, it remained in the nucleoplasm (lower panel). Images are representative of the GFP protein localization pattern during *P. infestans* infection. Bar =5 μm. **(C)** Data represent the number of CWC15-positive speckles during *P. infestans* infection. Bars represent the mean ± SD of 10 replicates from three independent experiments. ns=no significant difference and three asterisks denote a statistical difference at P<0.01 using two-sided Welch’s t-test.

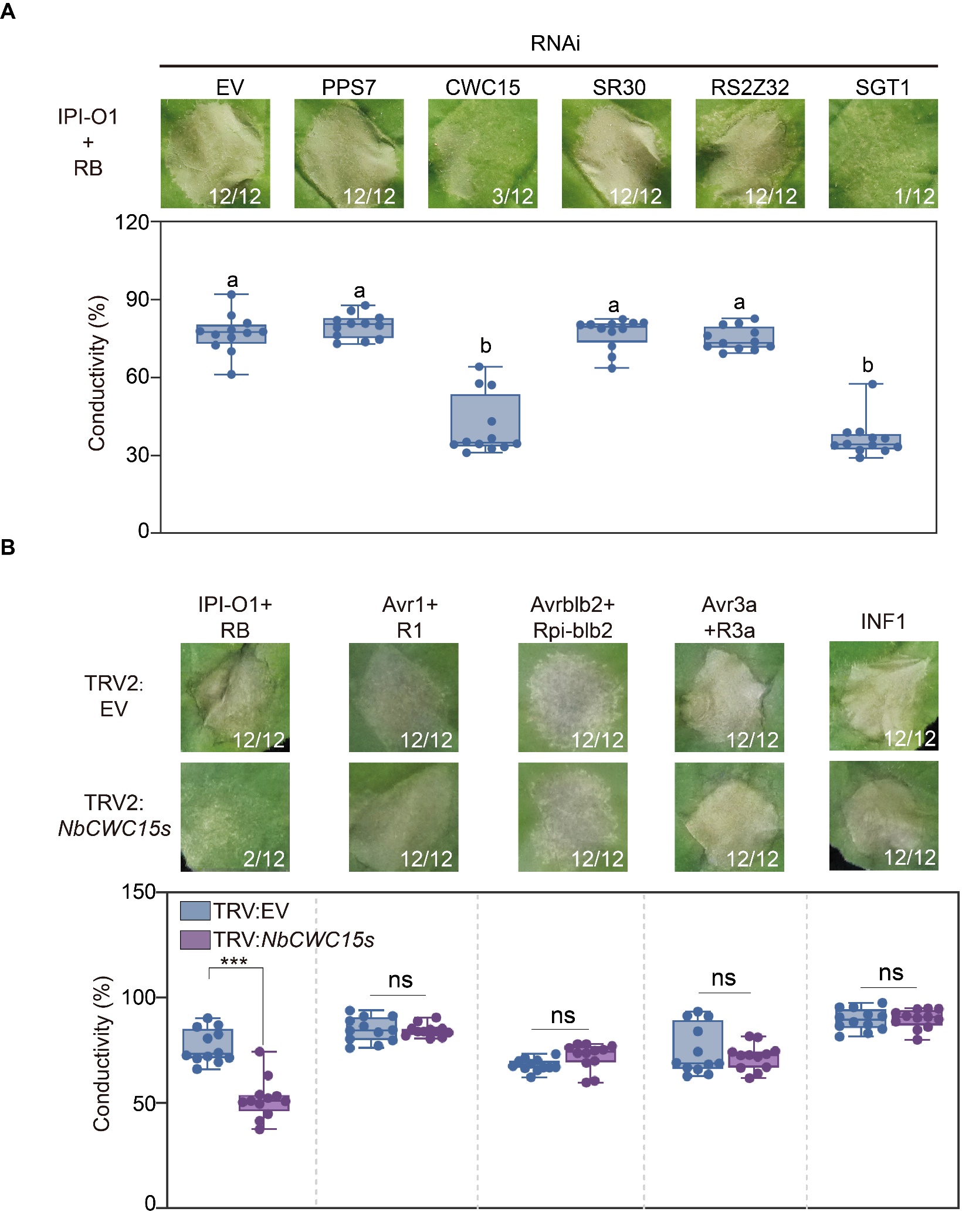

**Supplemental Figure S9**. **Splicing factor CWC15 is specifically involved in *RB*-mediated hypersensitive response.
(A)** Silencing of *CWC15* leads to loss of *RB*-mediated HR. IPI-O1 was co-expressed with the indicated RNAi constructs in *RB* transgenic *N. benthamiana*. Numbers represent the number of hypersensitive cell death responses compared to all infiltration points. Infiltrated areas were photographed 4 dpi. Electrolyte leakage was monitored by conductivity measurements following the same treatments (bottom). Data represent the mean ± SD from 12 replicates. Different letters designate a significant difference from the empty vector (EV) control at P<0.001 using one-way ANOVA with Tukey’s HSD test. **(B)** The designated Avr/R gene combinations were co-agroinfiltrated in *N. benthamiana* leaves following silencing of *CWC15*. The HR phenotype was photographed at 4 dpi. The numbers represent incidence ratios of cell death responses (upper). Below shows the results of electrolyte leakage assays of infiltrated areas. Data represent the mean ± SD from 12 replicates. ns = no significant difference and three asterisks denote a statistical difference at P<0.001 using two-way ANOVA with Tukey’s test.

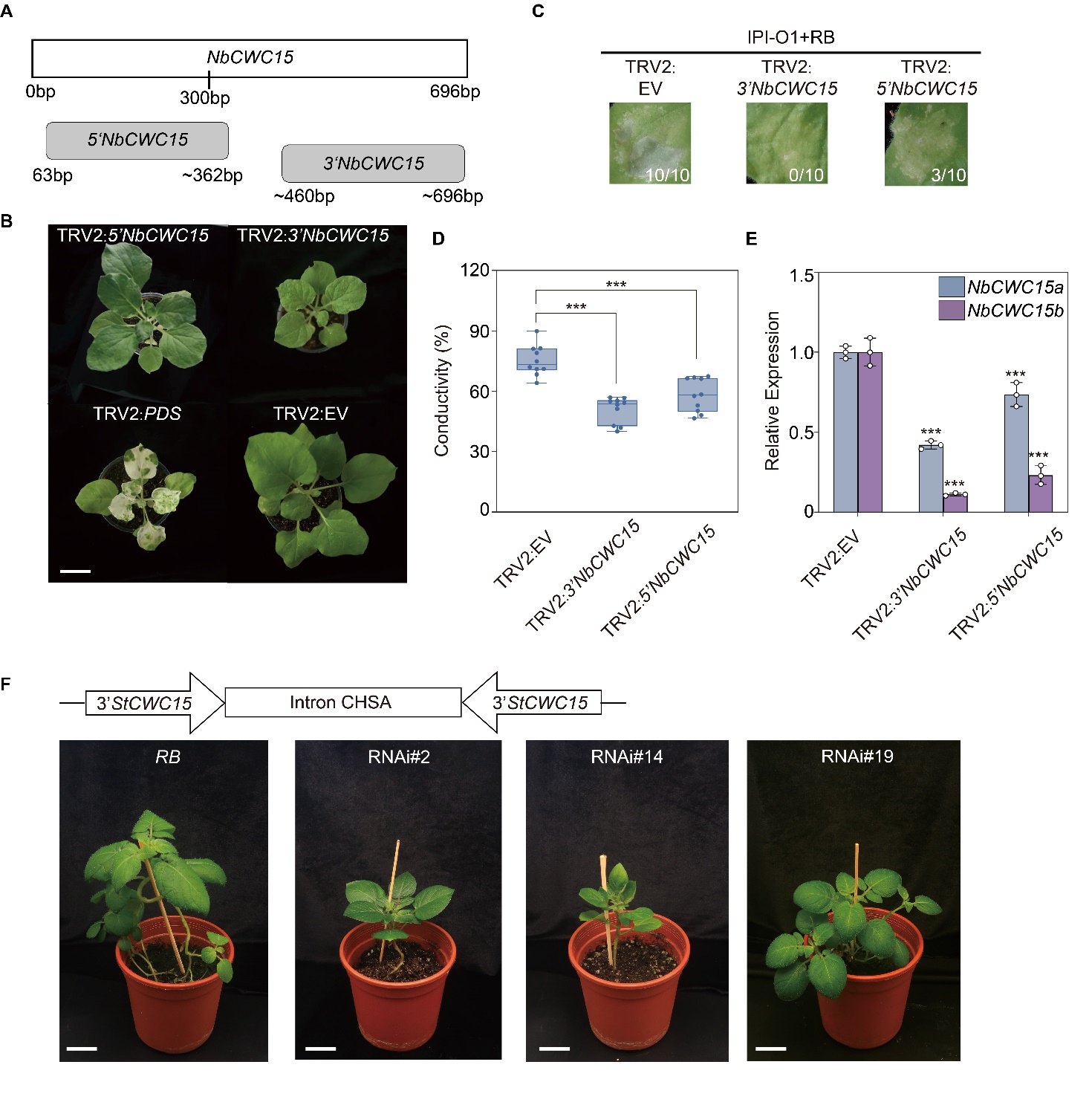

**Supplemental Figure S10. *CWC15* silencing in *N. benthamiana* by VIGS showed deficiency in growth and immunity.
(A)** Schematic diagram showing the fragments used in VIGS constructs. **(B)** The morphology of *NbCWC15* silencing in *N. benthamiana*. Two-week-old *N. benthamiana* plants were infiltrated with agrobacterium carrying TRV: EV, TRV: 5’*NbCWC15*, TRV: 3’*NbCWC15* and TRV: *PDS*, respectively. The photo was taken 3 weeks after infiltration. Bar = 3 cm. **(C)** 3’ *NbCWC15* or 5’ *NbCWC15* silencing has disturbed the recognition of IPI-O1 by RB. Hypersensitive cell death phenotypes were photographed at 4 dpi. The numbers represent the number of hypersensitive cell death responses compared to all infiltration points. **(D)** Electrolyte leakage from the infiltrated leaf discs was measured as a percentage of leakage from boiled discs at 4 dpi. Data represent the mean ± SD from 10 replicates. Three asterisks denote a statistical difference at P<0.001 using a one-way ANOVA with Tukey’s test. **(E)** Silencing efficiency of *NbCWC15* in *N. benthamiana*. Bars represent mean ± SD of three independent replicates. Two asterisks denote a statistical difference at P<0.001 using two-way ANOVA with Tukey’s test. **(F)** Above the photos is a schematic representation of the construct used for the *StCWC15* stable silencing by potato transformation. The photos show plant morphology of *RB* transgenic potato and *RB-StCWC15* RNAi transgenic potato lines following 30 days of growth in the greenhouse. Bar=3cm.

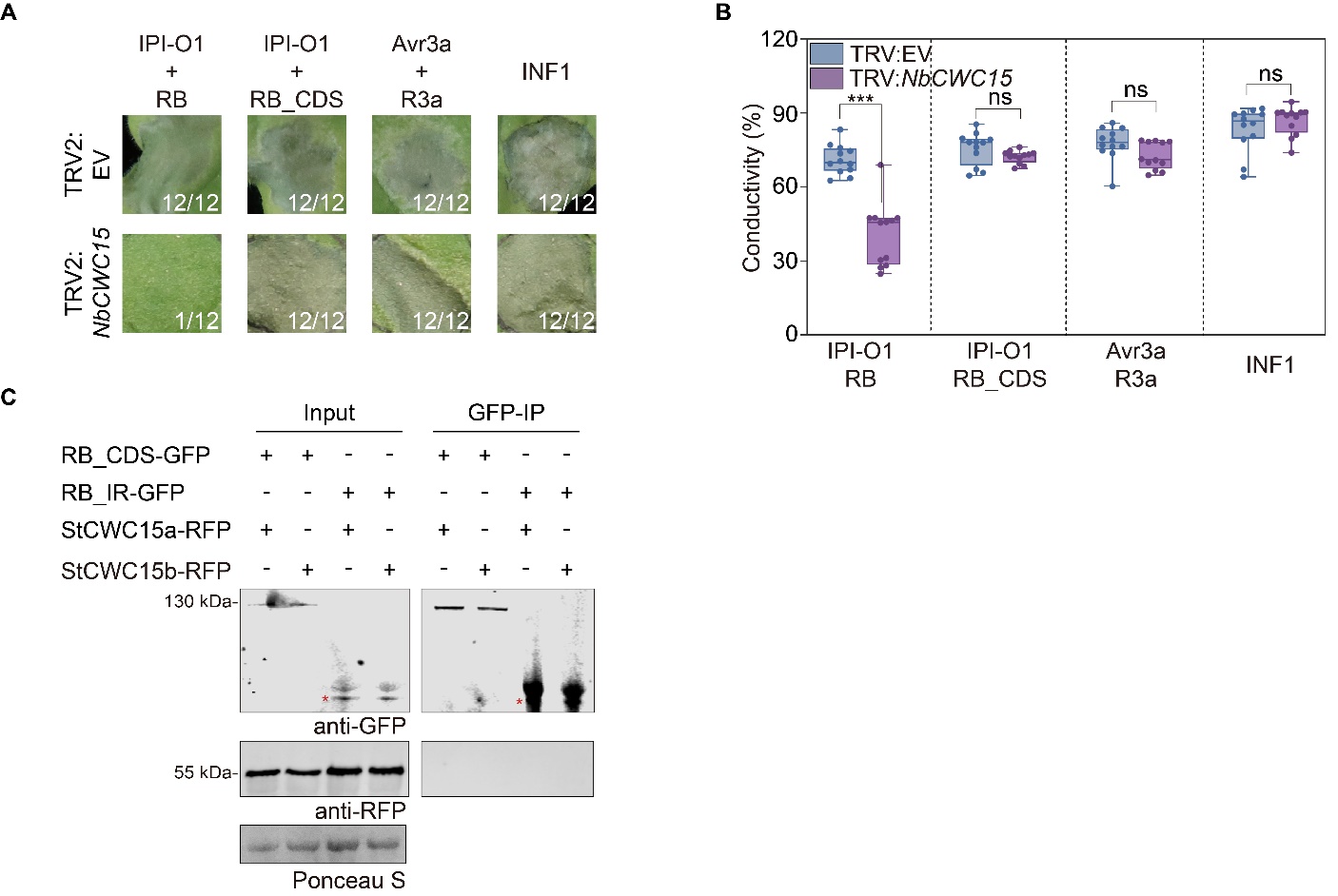

**Supplemental Figure S11. *CWC15* silencing has no effect on *RB_CDS* recognition of IPI-O1.
(A)** *CWC15* silencing impairs RB/IPI-O1 induced HR, but not RB_CDS/IPI-O1, R3a/Avr3a, or INF1. R gene/effector combinations and INF1 were agroinfiltrated into *NbCWC15*-silenced *N. benthamiana* leaves. Photos of HR phenotypes were taken at 4 dpi. The numbers represent exact number of hypersensitive cell death responses compared to all infiltration points. **(B)** Quantification of cell death by measuring electrolyte leakage at 4 dpi. Data represent the mean ± SD from 12 replicates. ns=no significant difference and three asterisks denote a statistical difference at P<0.001 using a two-way ANOVA with Tukey’s test. **(C)** RB_IR and RB_CDS do not interact with StCWC15a/b in vivo by co-IP. Immunoprecipitation (IP) of RB_CDS-GFP and RB_IR-GFP with StCWC15a-RFP, StCWC15b-RFP following agroinfiltration in *N. benthamiana* leaves. Expression of constructs in the leaves is indicated by +. Ponceau S staining was used to show levels of protein loading.

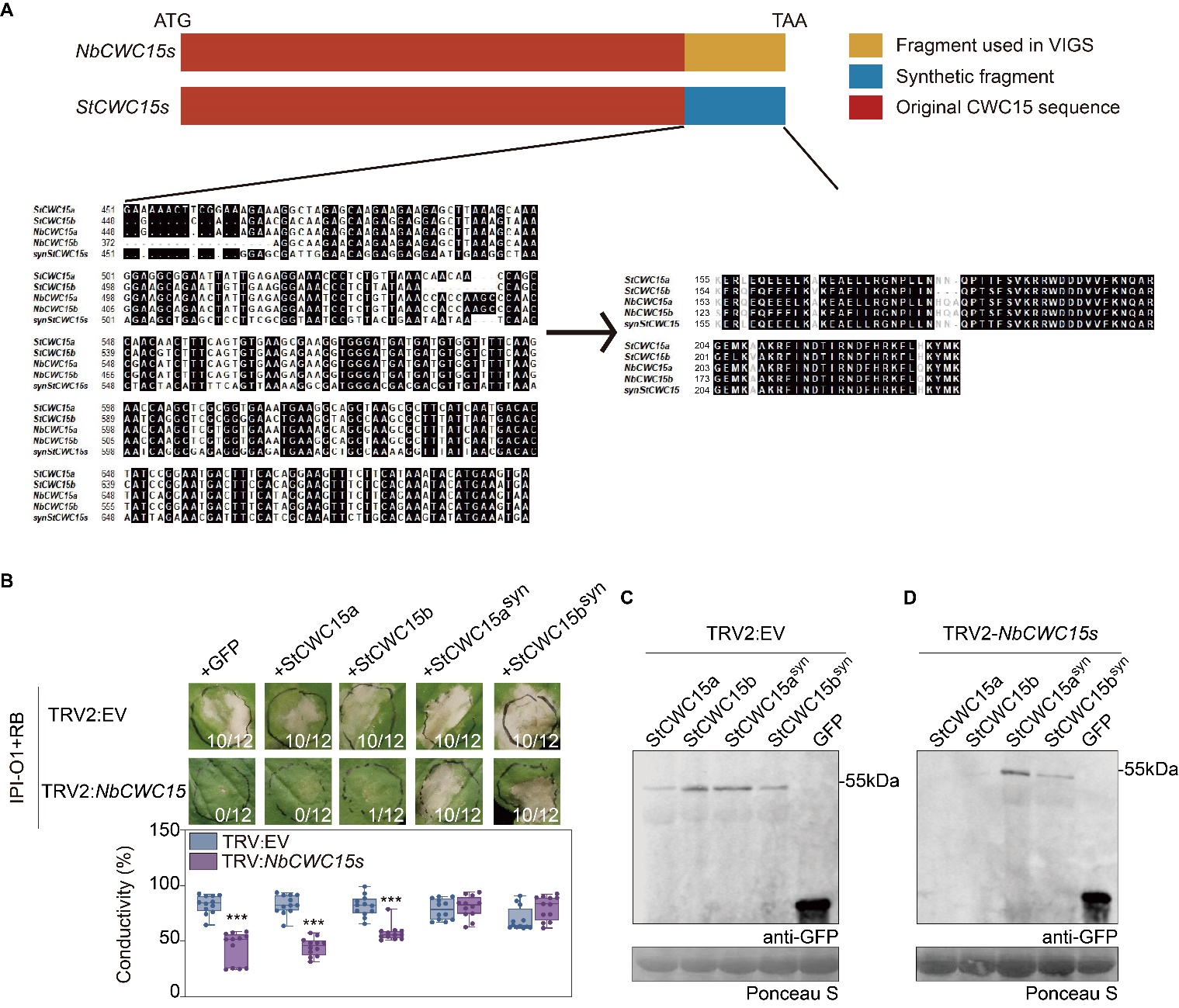

**Supplemental Figure S12. StCWC15 synthetic sequences complement *CWC15* silencing causing *RB*-mediated immunity deficient.
(A)** Schematic representation of VIGS and complementation design. The region from 460-696bp of *NbCWC15* was cloned into VIGS construct TRV2-3’*NbCWC15* for silencing. Synonymous substitutions were introduced into the synthetic *StCWC15* (*StCWC15^syn^*) without changing the protein sequence. The nucleotide and protein sequence alignments indicate the synonymous changes in the synthetic variant. **(B)** Expression of silencing-resistant synthetic *StCWC15* (*StCWC15a^syn^/ StCWC15b^syn^*) rescues *RB*–mediated HR in *NbCWC15*-silenced *N. benthamiana* plants. GFP and StCWC15a and StCWC15b were used as controls. Below shows results of electrolyte leakage assays of infiltrated areas at 4 dpi. The numbers represent the number of hypersensitive cell death responses compared to all infiltration points. Data represent the mean ± SD from 12 replicates. Three asterisks denote a statistical difference from the control at P<0.001 using a two-way ANOVA with Tukey’s test. **(C-D)** Immunoblot analysis of StCWC15a/b, StCWC15a^syn^/StCWC15b^syn^ in EV (C) and *CWC15*-silenced (D) plants. C-terminal GFP-tagged StCWC15 variants were transiently expressed in VIGS control and *CWC15*-silenced *N. benthamiana.* Protein loading was visualized by Ponceau S staining

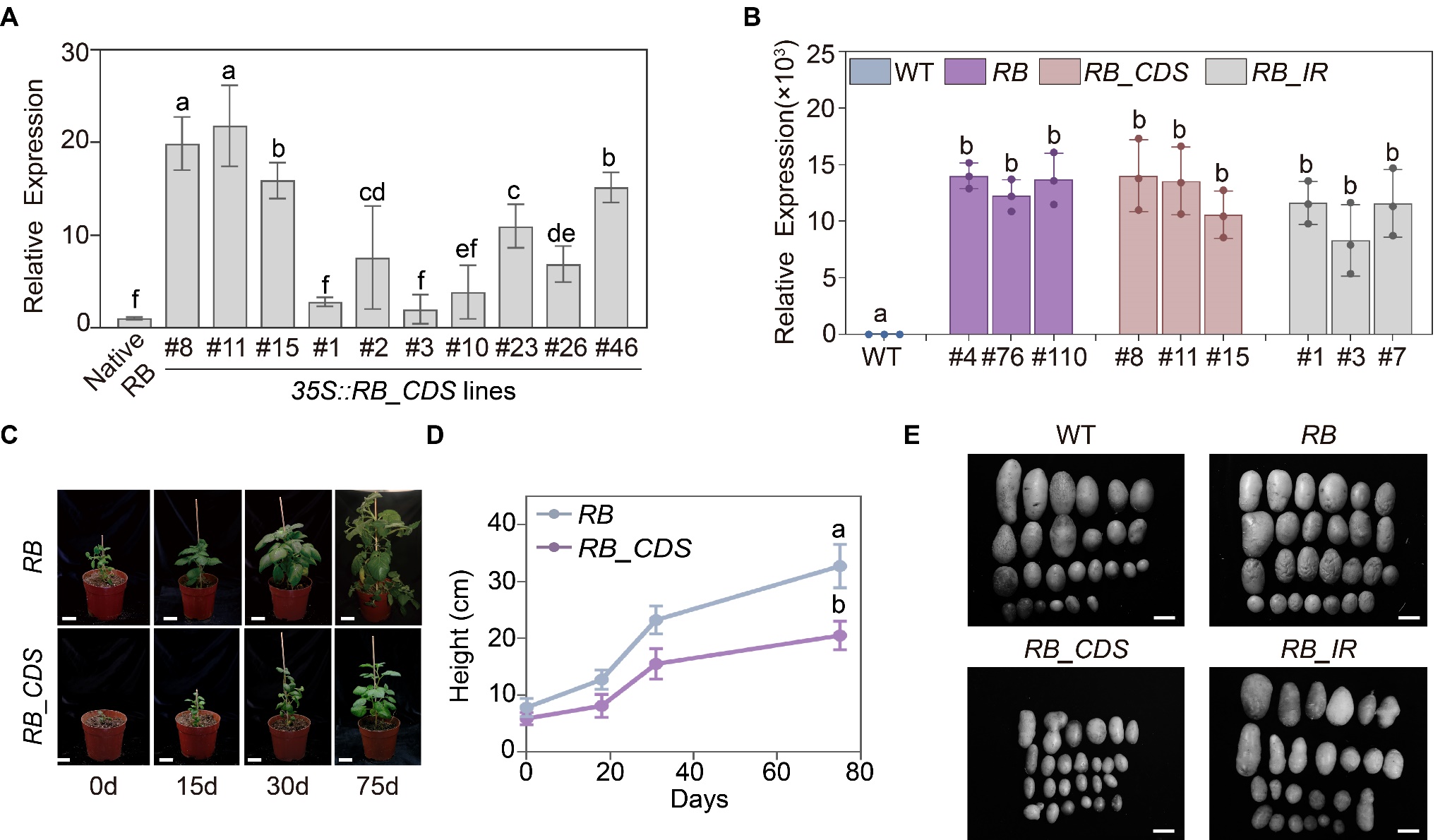

**Supplemental Figure S13**. **Overexpression of *RB_CDS* in potato causes growth abnormalities.**  **(A)** Expression level of *RB_CDS* in different *RB_CDS* transgenic potato lines. Bars represent the mean ± SD from three independent replicates. Different letters denote a significance difference at P<0.05 using one-way ANOVA with Tukey’s HSD test. **(B**) Expression level of *RB* in different *RB* transgenic potato lines. Bars represent the mean ± SD from three independent replicates. Different letters denote a significance difference at P<0.01 using one-way ANOVA with Tukey’s HSD test. **(C)** Photos of cv. Favorita expressing wildtype *RB* (Favorita^RB^) or *RB_CDS* (Favorita^RB_CDS^) from initial transplanting to 75 days after transplanting. Bar = 3cm. **(D)** The height of the transgenic plants graphed over time. Data points combine measurements from 5 individual transgenic plants. Data represent the mean ± SD from 3 replicates. Statistical differences among the samples were analyzed with one-tailed Student’s t test (P<0.05). **(E)** Tubers of Favorita (WT) and transgenic plants. Tubers were harvested and photographed 8 to 10 weeks after tissue culture plants were transplanted. Bar=1 cm.

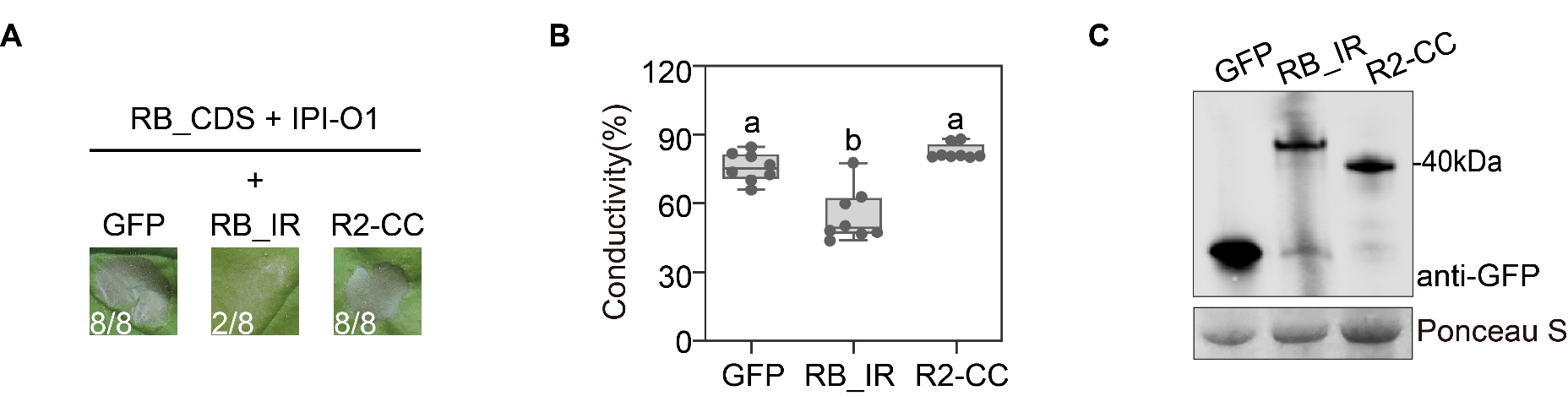

**Supplemental Figure S14**. **Overexpression RB_IR suppressed HR induced by IPI-O1 and RB_CDS.**

**(A)** IPI-O1 was expressed with the indicated proteins in *RB_CDS* transgenic *N. benthamiana*. Numbers represent the number of hypersensitive cell death responses compared to all infiltration points. Infiltrated areas were photographed 4 dpi. **(B)** Electrolyte leakage was monitored by conductivity measurements following the same treatments in (E). Data represent the mean ± SD from 8 replicates. Different letters above bars denote a significant difference at P<0.001 using one-way ANOVA with Tukey’s HSD test. **(C)** Immunoblot analysis of protein expression. Expression of GFP-RB_IR and GFP-R2-CC were confirmed by protein blotting using an anti-GFP antibody. Protein loading is visualized by Ponceau S staining.
